# Unsupervised learning of multi-omics data enables disease risk prediction in the UK Biobank

**DOI:** 10.1101/2025.10.02.679853

**Authors:** Chiara Rohrer, Justus F. Gräf, Marc Pielies Avelli, Ricardo Hernandez Medina, Henry Webel, Kirstine Ravn, Simon Rasmussen

**Affiliations:** Department of Biosystems Science and Engineering, ETH Zürich, Zürich, Switzerland; Novo Nordisk Foundation Center for Basic Metabolic Research, University of Copenhagen, Copenhagen, Denmark; Novo Nordisk Foundation Center for Biosustainability, Technical University of Denmark, Lyngby, Denmark

**Keywords:** VAE, multi-omics data, dataset properties, disease risk prediction, bias, feature importance

## Abstract

The size and complexity of biomedical datasets continue to grow, driving the development of methods that reduce dimensionality while preserving biological signals. Yet, when deep learning is applied to such data, the impact of preprocessing choices and dataset properties on model behavior is often overlooked. Here, we applied our framework Multi-Omics Variational autoEncoder (MOVE) to multiomics data from 452,026 UK Biobank participants, aiming to both evaluate the power of the learned representations for disease risk prediction and critically analyze how non-biological factors, like dataset properties and preprocessing decisions, can shape and influence the results. We show that reducing the dimensionality of the data by a factor of 80 still yields comparable prediction performance across 15 different diseases. We further demonstrate how dataset properties and preprocessing choices impact the model performance, latent representation and downstream results, and our findings strongly underline the need for thorough analysis and understanding of a model’s behavior before drawing conclusions from its results.

## Introduction

Large biomedical datasets are an invaluable resource for the study of disease prevention, onset and progression. Such datasets can offer insights into the relations established between biomarkers and their role in the development of disease ^1–3^. The identification of measurable factors associated with a future disease risk can enable the stratification of individuals into risk groups^4^, thus helping with prevention and early diagnosis. This stratification can also be extended to patients^5^, who can benefit from tailored treatments^6^. The past decades have brought many of these large-scale initiatives, including the UK Biobank^7^ and the All of Us research program^8^, which focus on representing individuals and follow them prospectively. Another example is the Adult Genotype Tissue Expression (GTEx) project^9^, which is devoted to the analysis of multiple tissues from deceased individuals.

These high-dimensional datasets often contain multiple types of biological data. We refer to these as “multi-omics”, a term that encompasses genomics, transcriptomics, proteomics, and metabolomics among others. Despite the large number of measured variables, many of them are often associated or correlated, and hence the datasets are somewhat redundant. Several approaches to integrate multi-omics data have been developed^10,11^, exploiting such redundancy to reduce the dimensionality of the data without losing biologically relevant information. However, biobank-scale datasets often contain missing values, both across features and patients: a given individual might have very few measurements, or a variable might be measured in only a few individuals. Both cases can pose problems when trying to integrate the data, and while a straightforward option is to remove samples or features with high missingness rates^12^, valuable information might be discarded in the process and more satisfactory solutions are required.

Deep learning (DL) based methods have emerged as a promising avenue for solving these challenges. A popular approach to reduce the dimensionality of large datasets in an unsupervised manner is the use of Variational AutoEncoders (VAE). These models are trained to learn a lower dimensional representation of the data, which is called latent, while reconstructing the input as precisely as possible^13^. VAEs have been used to compress electrocardiogram (ECG) data^14^, imaging data^15^, single-omics datasets^16,17^ and multi-omics datasets^18^. In addition, several VAE frameworks have been developed to specifically address the characteristics of biomedical datasets, such as our framework, the “Multi-Omics Variational autoEncoder” (MOVE)^19^. MOVE can be trained on data with missing values, integrate multi-modal measurements to more meaningful lower-dimensional representations, and identify drug-omics or cross-omics associations through *in silico* perturbations.

When using DL based models such as VAEs in a clinical setting, it is desirable to understand the model’s behavior. This ensures transparency, reliability, and trust in these methods while preventing potentially fatal errors due to model biases^20^. Therefore, the development of models using clinical data is often accompanied by attempts to understand the model’s decision-making process^21,22^. These can be in the form of Shapley additive explanations (SHAP)^23^, or similar feature attribution mechanisms where calculated feature importances give insights into what a model learned and used for the prediction tasks.

The task of predicting disease risk can in fact be framed in several ways. It is possible to fit a regression model to infer the time to diagnosis, or a classification model to predict the occurrence of the disease. However, both approaches come with limitations due to the right-censored format of the data. In regression models, only individuals with diagnoses can be used, while for the classification models, all information regarding time is lost. Survival models emerge as a promising alternative when modelling individual disease risks based on the provided data. Models such as the Cox proportional hazards model^24^ have been used on proteomics or metabolomics data^25–27^. There is also a growing number of DL based survival models^28^ that can be trained on higher dimensional datasets compared to standard survival models^29^.

Here, we integrated UK Biobank multi-omics data including blood, urine, physical measurements, as well as metabolomics, proteomics, genomics, and seroprevalence data of infectious diseases using the DL framework MOVE. The predictive power of the learned disease-agnostic embeddings matched that of a disease-dependent subset of the input features while compressing the input dimensionality roughly 80-fold. The analysis of survival models showed that MOVE learned to rely on biologically meaningful features and that the top predictive features aligned with standard diagnosis procedures. However, data specific properties of the modalities, as well as the preprocessing methods and hyperparameter choices influenced the organization of the latent space. These in turn influenced the model’s ability to reconstruct the compressed samples and to impute missing values. Altogether our results showcase the intricate interplay of biological and non-biological signals in the data and the need to disentangle their effects when working with large biomedical datasets.

## Results

### UK Biobank wide multi-omics data integration and survival analysis using MOVE

We first applied MOVE^19^ to perform unsupervised integration of the biological modalities in the UK Biobank. The obtained latent representations of the individuals were then used to train and evaluate disease-specific survival models (**Figure 1a**). Our analysis included 452,026 individuals across nine modalities: phenotype-associated genotype data (“genomics”), protein Quantitative Trait Loci (pQTL), affinity-based proteomics, metabolomics, blood biochemistry, blood cell counts, physical measurements, urine measurements, and infectious disease seroprevalences. In total, we included 10,277 features across all modalities. Of these features, 7,089 were categorical while the remaining 3,188 were continuous. Most of the features came from the genomics dataset (3,943, 38.4%), while the smallest dataset described urine biomarkers (4 features, 0.04%) (**Figure 1b**). Data completeness varied markedly across modalities, features, and individuals. Overall completeness of the full dataset was 73.2%, with average completeness <10% for proteomics and infectious disease seroprevalence data,>90% for physical measurements and 100% for the single nucleotide polymorphism (SNP)-based modalities (genomics and pQTLs) (**Figure 1c**). Most individuals had data for approximately 70% of the features, while few were in the range between 80% and 100% (**Figure 1d**). MOVE was trained using 90% of the individuals (406,823) to perform nonlinear dimensionality reduction, and multiple types of disease-specific survival models were built on top of the obtained latent representation. The remaining 10% of the individuals was used as a test set for evaluation of the survival models. Finally, we analyzed MOVE’s data reconstructions and latent representations and compared the performances of the different survival models (**Figure 1a**).

**Figure 1:**
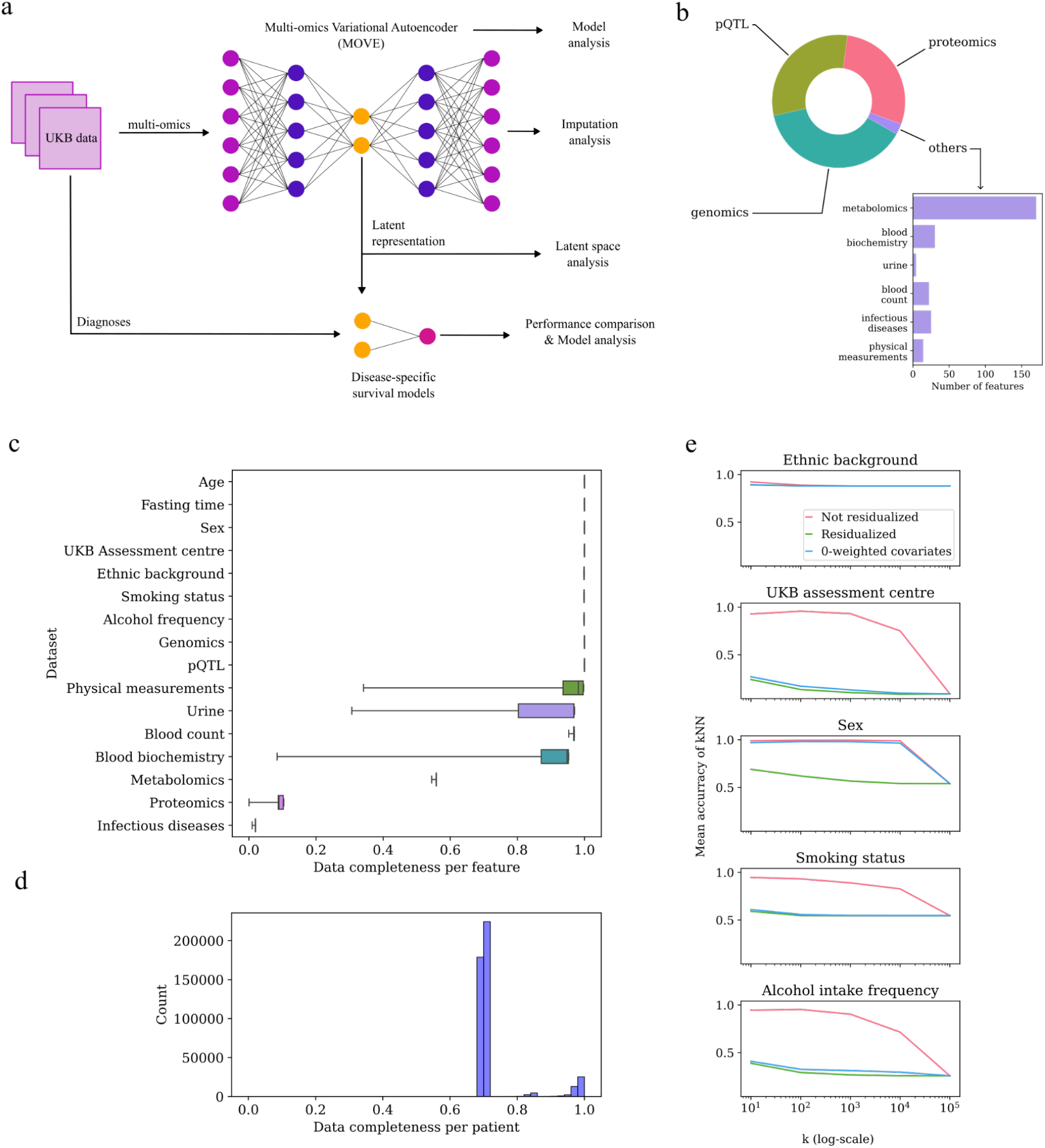
Analysis overview, data characteristics and effect of residualization. **a**) Visual abstract of models and subsequent analysis. A multi-omics dataset was learned by MOVE (top), latent representation was used by different types of survival models for disease risk prediction (bottom). Models and obtained representations were analyzed and compared (right). **b**) Distribution of features per modality in the residualized dataset. **c**) Data completeness per features, grouped by modality and including covariates. Each box displays the three quartiles and the whiskers mark points in the corresponding 1.5 interquartile ranges. **d**) Data completeness by patient, including only features from the nine used modalities. **e**) The effect of dataset residualization on clustering by categorical covariates is displayed by quantifying clustering in the latent space. For each categorical covariate, the mean accuracy of the kNN classifier is displayed for three differently preprocessed datasets (red: not residualized, green: residualized, blue: 0-weighted, non-residualized) and different values of k. Note that in some cases, lines might lie on top of each other and are therefore not fully visible.

### Residualization of covariates reduced clustering of samples in the latent space

When integrating the data using MOVE, we observed that the inclusion of categorical covariates as inputs to the model led samples to cluster in the latent space (**Supplementary Figure 1a**). To understand how input data preprocessing affected the latent space, we trained three versions of the MOVE model, varying the inclusion of covariates, residualization, and the contributions of different data modalities to the loss function. The degree of clustering was quantified using the mean accuracy of a k-nearest neighbour (kNN) classifier across different neighborhood sizes (k = 10 to 99,999). High kNN accuracy indicated strong clustering by categorical features, while values close to the majority class indicated well-mixed representations.

Residualization of continuous features using the covariates substantially reduced the covariate effects on the latent space. The mean decrease in kNN accuracy across all *k* when comparing non-residualized to residualized data was 0.601 for the UK Biobank assessment centre, 0.008 for ethnicity, 0.311 for sex, 0.273 for smoking status, and 0.463 for alcohol intake frequency. Alternatively, including covariates as inputs to the model and assigning them zero weight in the loss function reduced clustering almost as well as the residualized dataset for some covariates (**Supplementary Figure 1b**). This approach closely approximated residualization for UK Biobank assessment centre (mean difference: 0.021) but was less effective for sex (mean difference: 0.296) (**Figure 1e**). Notably, clustering by ethnicity was observed regardless of the method used. We hypothesized that the persistent ethnicity clustering was caused by the inclusion of categorical genotype data (genomics and pQTLs), which grouped the individuals by genetic similarity. To address this, we encoded the genotype data as continuous variables, allowing them to be residualized. While this successfully reduced the influence of ethnicity, it also reduced the ability of the VAE to reconstruct the samples (mean reconstruction accuracy: 0.806 vs. 0.890) (**Supplementary Figures 1d, 2e, 2c**).

As these findings indicated a tradeoff between reconstruction accuracy and suppression of covariate-driven clustering, we proceeded with the latent space generated from the residualized dataset while still treating genotypes as categorical variables. This promoted a rather homogenous latent representation while maintaining good reconstruction metrics.

### Adjusting the relative importance of each data modality improved reconstruction accuracy

Initially, the contribution of each data modality to the loss was weighted equally. We noted that this approach would lead modalities with high missingness (e.g. proteomics) to strongly influence the sample arrangement in the latent space. This was manifested by the model grouping individuals with available proteomics data in the same region of the latent space (**Supplementary Figure 2b**), a behavior that suggested that the model was not effectively learning how to represent features with many missing values.

Aiming to compensate for this effect, we adjusted the loss calculation by weighting the contribution of each data modality inversely to the degree of data missingness (**Methods**). Although this did not reduce the clustering based on data missingness (**Supplementary Figure 2d**), it improved reconstruction metrics for the sparse modalities. Cosine similarity between the input and reconstruction of the features, treating each modality as a vector per individual, increased by 0.098 for proteomics, 0.014 for metabolomics, 0.003 for urine measurements and 0.110 for infectious disease prevalence. For the remaining datasets, decreases in performance were minimal, with less than 0.015 and 0.009 in accuracy and cosine similarity, respectively (**Supplementary Figures 2a, 2c**). Overall, the reweighted model achieved a mean cosine similarity of 0.968 for continuous modalities and a mean accuracy of 0.709 for categorical modalities. Finally, the proteomics data had the lowest mean cosine similarity (0.648), while the urine biomarkers had the most accurate reconstruction on average (0.988) (**Supplementary Figure 2c**). The improved reconstruction metrics led us to proceed with the model where the loss contributions of each data modality were adjusted.

### Datasets with high data completeness dominated the latent representations

To understand the structure of the latent space learned by MOVE and identify which features influenced it most, we calculated feature attributions for the encoder and decoder networks (**Methods**). Using the mean of the absolute attribution values across 5,000 randomly sampled individuals, we found that the latent representation was primarily influenced by blood biochemistry, blood cell counts, physical measurements, and urine measurements (**Figure 2a, left**). When aggregating encoder attributions by modalities, the genomics and pQTL datasets had the highest contribution (**Figure 2a, right**). Notably, features with low data completeness contributed less to the latent representations than features with higher data completeness. This pattern was supported by a strong positive correlation between a feature’s encoder importance and its data completeness (Pearson r = 0.801) (**Figure 2b**). This suggests that MOVE preferentially relied on features with high completeness, likely because sparse features were harder to reconstruct. On the other hand, individual feature importance tended to be lower in datasets with many highly correlated features (e.g. genomics). We interpret this as the model distributing importance across redundant features, thereby reducing the per-feature attribution while still encoding relevant biological signals. These results suggested that the composition of the latent space was shaped by both data completeness and the degree of feature redundancy within each modality, indicating how non-biological data properties like feature availability, feature completeness and redundancy can influence what the model learns.

**Figure 2:**
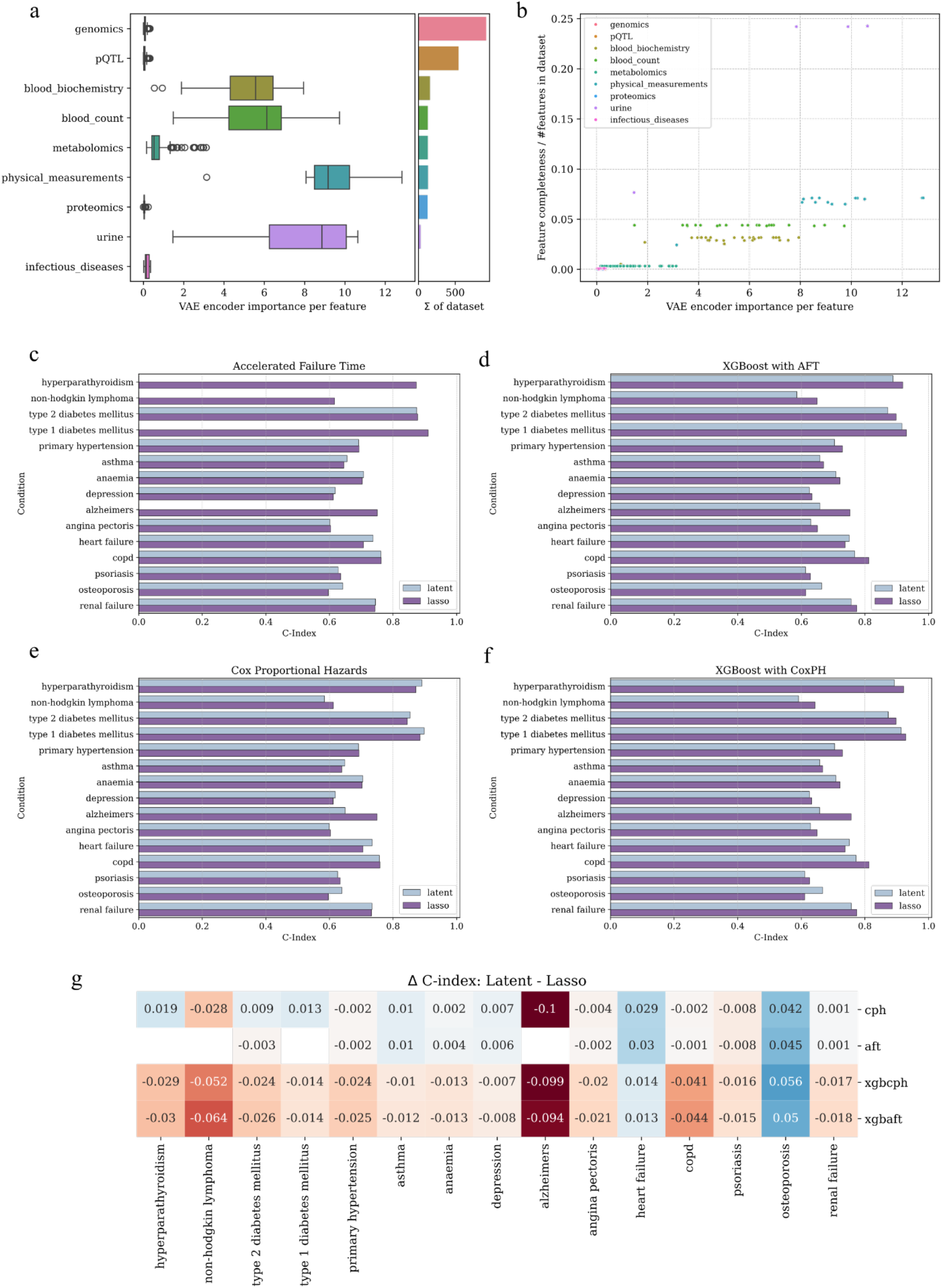
Results of MOVE and survival models. **a**) Feature importances for encoder grouped by modality (left) and sum over all features per modality (right). **b**) Correlation between encoder importance and fraction of data completeness and number of features in corresponding modality, colored by modality. **c-f**) Performances of all survival models, grouped by diseases. Different colors indicate different input features: “latent” for the disease-independent, 128-dimensional latent space of MOVE, “lasso” for the disease-specific lasso-selected feature subset. **g**) Differences in model performances when using latent representation and disease-specific lasso-selected feature subset. Missing values indicate that one model failed during training. Abbreviations: “cph” = Cox proportional hazards, “aft” = Accelerated failure time, “xgbcph” = XGBoost with Cox-loss, “xgbaft” = XGBoost with AFT-loss.

### Latent representations match disease-specific features in survival model performance

To evaluate whether the latent representations learned by MOVE retained clinically relevant information towards disease incidence prediction, we compared several survival models trained on those multi-modal latent representations with models trained on disease-specific feature subsets. For each of 15 selected diseases, spanning a variety of conditions, we trained four types of survival models to predict 10-year incidence: Cox proportional hazards models^24^, Accelerated Failure Time (AFT) models^30^ as well as XGBoost-based models^31^.

The predictive performance of the survival models trained on latent representations was comparable to that of the models trained on lasso-selected input features. The mean difference in concordance (C-index) between the two approaches was -0.0008 for the Cox models, and 0.0073 for the AFT models -excluding the four AFT models that failed during training (**Figures 2c, 2e**). For the XGBoost approach, the lasso-selected features performed slightly better than the model trained on the latent representation, with mean improvements of +0.020 (Cox-loss) and +0.021 (AFT-loss) (**Figures 2d, 2f**). Survival model performance varied more pronouncedly depending on the predicted disease. Hyperparathyroidism, type 1 diabetes, and type 2 diabetes showed the highest predictive performance across all four models, with C-indices exceeding 0.8 for the successfully trained models. Notably, the models based on the latent representation outperformed those using lasso-selected features for osteoporosis and heart failure. This was the case across all four survival model types with average C-index improvements of 0.048 and 0.022, respectively (**Figure 2g**).

Overall, our results demonstrate that disease-independent, compressed latent representations obtained using the framework MOVE preserve biological signals that can be used for disease risk prediction. Models trained on such representations yielded performances comparable to those trained on disease-specific feature subsets, both when using traditional and gradient-boosted survival models.

### MOVE initialization had a minor effect on survival model performance

Aiming to quantify the impact of MOVE’s initialization conditions on the downstream survival model performance, we repeated the entire modeling pipeline multiple times using different random seeds (**Methods**). While the survival models themselves produced consistent results when retrained from fixed inputs, variation in the latent representations from different MOVE initializations led to small but measurable differences in C-index values (**Supplementary Figure 3**). The magnitude of this variation was both model and disease-dependent. The largest difference in C-indices was observed for type 1 diabetes mellitus (ΔC-index = 0.049) trained using the XGBoost model with Cox loss, whereas the smallest was observed for primary hypertension using the Cox proportional hazards model (ΔC-index = 0.004). Therefore, while MOVE introduced slight variations in the latent representations each time it was trained, the effect on downstream predictive performance was generally minor and disease-dependent.

### Predictive features were aligned with clinical diagnosis guidelines

To investigate the factors driving the strong predictive performance for type 1 diabetes mellitus, hyperparathyroidism, and type 2 diabetes mellitus, we calculated SHapley Additive exPlanations (SHAP)-values^23^ for the lasso-selected input features. We found that the top predictive features were closely aligned with established clinical diagnostic criteria^32,33^. Calcium, phosphate, and albumin were the most predictive features for hyperparathyroidism. HbA1c and blood glucose were the drivers of type 1 diabetes predictions. Lastly, HbA1c, waist circumference and blood glucose were prioritized when predicting type 2 diabetes. These findings confirmed that survival models were able to base their predictions on biologically meaningful features and showed how explainability methods such as SHAP can aid in the interpretation of the results, including the validation of biological signals.

### The predictions of different diseases relied on different latent dimensions

Understanding what the survival models learned from the latent representations was not trivial, as the latent dimensions in a VAE do not necessarily carry any intrinsic meaning. To work in that direction, we first calculated SHAP-values for the XGBoost models and retrieved the coefficients for the Cox and AFT models. We noted that, while many latent dimensions do not have much influence on the prediction, the set of influential latent dimensions and their magnitude changed depending on the disease (**Supplementary Figure 4**). When summing feature importances across all latent dimensions per disease, we observed strong correlations between the sum of absolute feature importances and the C-indices (Pearson correlation between 0.73 and 0.92, p-values < 0.005). These results suggested that the performance of the survival models relied on the availability of predictive information encoded in the features learned by the VAE. The selected set of features might be more useful to predict certain diseases than others, which would also affect the corresponding latent representations of the samples. This would explain the difference in predictive performance across diseases for both types of input data. To assess which features ultimately informed the disease predictions from the latent space, we matrix-multiplied the feature importances of the survival model with the feature importances of the VAE encoder. We found that the most important features across all diseases came from the four datasets with the highest encoder feature importances, namely urine measurements, physical measurements, blood biochemistry and blood count. This highlighted the influence of the encoder on the feature importances for disease risk prediction (**Supplementary Figure 5**) and underlines how influential the initial VAE training is for downstream results in this workflow.

### Mapping single-omics datasets to the latent space does not improve survival model performance

Next, we wanted to evaluate MOVE’s ability to represent and reconstruct individuals for which only one data modality was available. We initially hypothesized that the pre-trained encoder might add information to the mono-modal data when mapping the patient’s representation to a shared latent space shaped by all data modalities. This could be tested by comparing the C-indices of the different disease prediction models, i.e. those trained on the latent space representations of the samples with those trained on the single-omics input datasets.

As shown in **Supplementary Tables 1-7**, mapping individuals to the latent space did not significantly improve survival model performance across all datasets, model types or diseases. Of note, there were often too few positive disease cases to get a reliable performance score. Out of the 46 (of 127) combinations of a disease and a mono-omics dataset where there at least 100 incidences in the test set, in 13 cases the best latent model outperformed the best model on the original (single-omics) or lasso-selected data. However, the mean improvement for these 13 cases was 0.016, while the maximum was 0.048 (prediction of angina pectoris using genomics).

We therefore conclude that the latent representations obtained using MOVE were not able to systematically enhance the survival model performance of single-omics datasets by mapping the patients to the shared latent space.

### Imputation quality depends on the properties of the dataset

We finally examined whether the pre-trained MOVE model could act as an imputation method for missing values in the dataset. The framework’s imputation power changed depending on the nature of the missingness, e.g. when randomly removing a fixed number of observations per feature or when removing full datasets at once (**Methods**).

The removal of 2,000 samples per feature at random led to some features being very well correlated with their original values (5.5% of the features had Pearson correlation greater than 0.8), but many had a correlation around 0. This behavior depended on the data modality, but there was generally a moderate trend showing better reconstruction for features with a larger standard deviation in their reconstruction (Pearson correlation of 0.562) (**Figure 3d**). Notably, the metabolomics dataset had the best mean Pearson correlation coefficient between original and imputed values across all features (0.949), while the mean across all datasets was 0.270 (**Figure 3b**).

**Figure 3:**
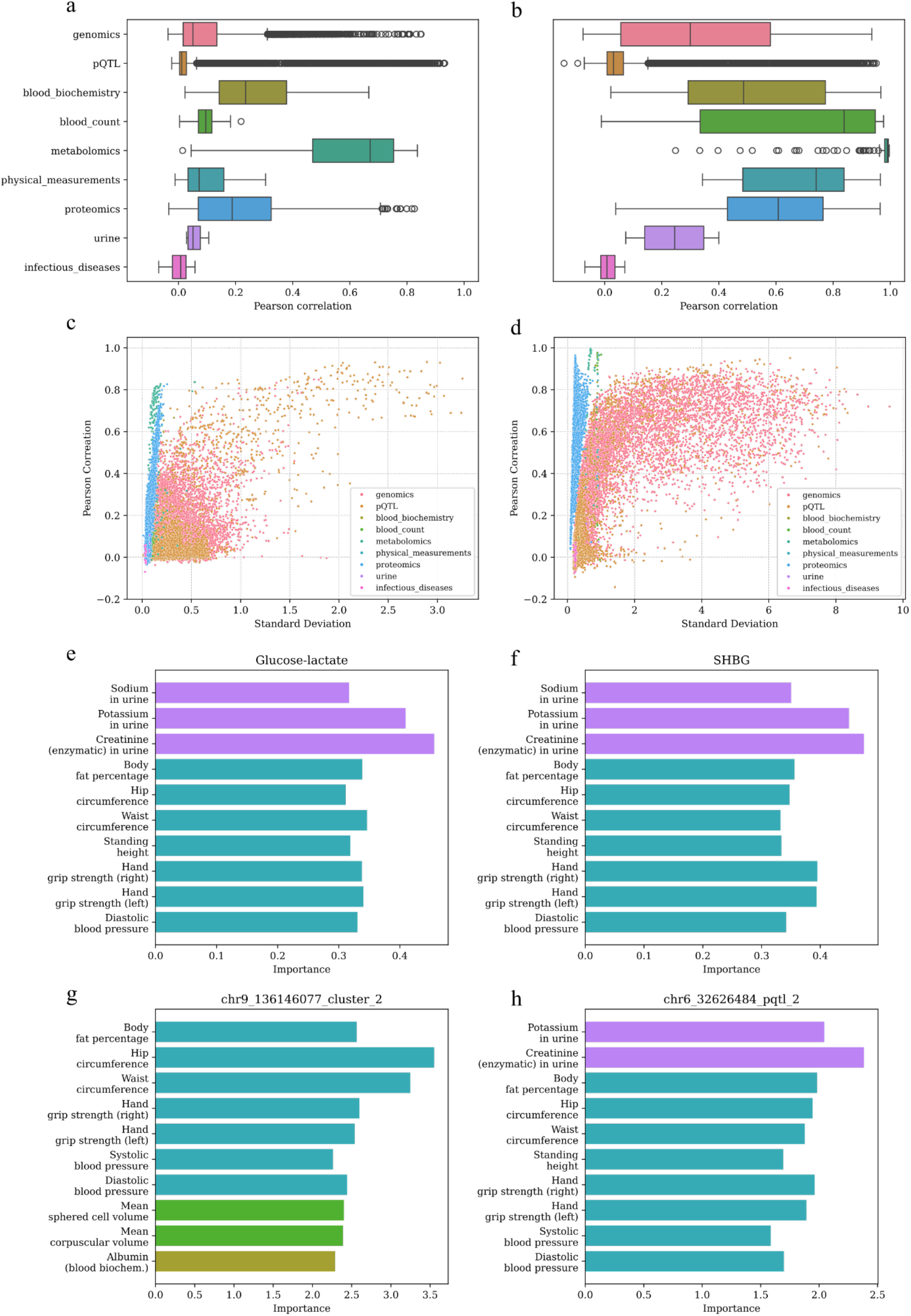
Analysis results of feature importances and imputation performance of MOVE. **a**) Imputation quality (measured by Pearson correlation) when single modalities were completely removed, grouped by modality. b) Imputation quality (measured by Pearson correlation) when 2,000 values were randomly removed per feature across all modalities, grouped by modality. **c**) Imputation quality (measured by Pearson correlation) and standard deviation of imputed features when single modalities were completely removed. **d**) Imputation quality (measured by Pearson correlation) and standard deviation of imputed features when 2,000 values were randomly removed per feature across all modalities. **e**)-**h**) Most important features for reconstruction for the best-reconstructed features of metabolomics (**e**), proteomics (**f**), genomics (**g**) and pQTL (**h**) given in absolute feature attributions and colored by modality.

Conversely, when removing one full dataset at a time we generally found that the Pearson correlation (0.351) between original and imputed values was worse than that of the first missingness scenario. Only 0.5% of the reconstructed features had a Pearson correlation with their original values greater than 0.8. Again, we observed a relation between the standard deviation of the reconstructed feature and the Pearson correlation between reconstructions (**Figure 3c**). Furthermore, the metabolomics dataset had the best mean Pearson correlation across all features (0.590) while the mean over all datasets was 0.099 (**Figure 3a**). This suggested that the model learned shared information across datasets, especially between the metabolomics dataset and the other eight. We then calculated the top 10 input features, by importance, for datasets with at least one feature with Pearson correlation above 0.8 (metabolomics, proteomics, genomics, pQTL). In all four cases, the most important features were found in the four datasets which also had the most influence on the latent space, as discussed previously (**Figures 3e-h**).

In summary, the ability of MOVE to reconstruct missing values depended not only on the pattern of data missingness in the dataset, but also on the ability of the model to connect features with shared information across different datasets. Additionally, well-reconstructed features seemed to be informed by datasets that had the highest influence on the latent representations of the samples, which in turn were, as previously shown, strongly driven by dataset properties like feature completeness.

## Discussion

In this study, we used the VAE-based framework MOVE to integrate and compress the multi-modal data in the UK Biobank dataset, a large multi-omics study including nearly half a million individuals. Aiming to predict disease risk for 15 different diseases, we trained a number of survival models on the compressed sample representations learned by the models. The information encoded in such disease-independent representations of the samples led the survival models to achieve comparable performances to those trained on a disease-relevant subset of input features. This is quite remarkable, as the VAE was forced to compress the data from 24,455 input features to only 128 dimensions in the latent space. This corresponds to a compression by a factor of 80 when considering that the original data had 10,277 features. As data compression generally leads to information loss, the observation that all models trained on the latent representation outperformed the models on the lasso-selected subset of original features for both heart failure and osteoporosis was unexpected, motivating further research to determine the underlying drivers of this result. In case the data integration truly facilitated disease risk prediction for some diseases, this would highlight the ability of MOVE to learn biologically meaningful representations. Alternatively, fine-tuning the VAE encoder after training to adapt the latent representation to less well-performing diseases might possibly enhance the performance of the survival models.

Despite their widespread applications in the field, traditional survival models and their use are hotly debated. The underlying assumptions of these models are not met when using high-dimensional clinical data, and yet it is not obvious nor globally accepted that they ought to be met at all^34,35^. Further research into the effects of these violated assumptions on model predictions would lead to an increased trust in the models when applied to clinical settings. The use of C-index as a survival performance metric can be biased^36^, but we note that this has little to no effect on our analysis within diseases, as we compare different models based on the same diagnosis data. This bias could nevertheless affect the performance comparison across diseases, as the number of incidences varies. An additional source of bias was the data used to define the time-to-diagnosis, as we only had access to data from hospital inpatients. Models might be biased if the disease was often diagnosed in a non-hospital setting.

To interpret the survival model predictions, we assayed feature importance attributions. High-scoring features in models feeding on the lasso-selected feature set generally agreed with common diagnosis guidelines^32,33^. However, this behavior was not associated with a higher prediction performance, since it was observed for diseases with a wide range of C-indexes. When studying the feature importances for the survival models trained on the latent representations, we mainly identified strong signals from the set of features driving the latent representations. We observed a strong relation between assigned feature importances, feature completeness and the number of features in the dataset. Such dependence on data completeness could be explained by the model giving a higher weight to more “reliable” features, while the dependence on the number of features in the dataset can be explained by the attributions being distributed across similar features. The latter was mostly relevant to SNP data, as one-hot encoding led to a high degree of redundancy in the genomics and pQTL datasets. These results demonstrate the strong influence of the encoder on downstream applications and remark the importance of understanding the factors driving the latent representations. We therefore see a strong need to critically question biological findings by analyzing both the properties which drive the used models output and the methods itself, especially with regards to assumptions and the potential violations thereof.

In addition to the disease risk predictions, we tested whether the pre-trained VAE could be used for other applications. We noted that the trained MOVE model could not enhance disease incidence prediction for single-omics datasets. Given its fully connected architecture, the encoder relied on all input features. As illustrated by our feature importance attribution analyses, the absence of key features heavily changed the positioning of the samples in latent space. This in turn shaped the subsequent reconstruction of the feature profiles at the decoder, which also included the missing data modalities. Furthermore, when using only samples from a single dataset, the number of incidence cases in the training and test sets were reduced in comparison to the full dataset. This probably affected both the survival model training and evaluation, as the number of incidence cases for certain diseases gets critically low for some data modalities. To avoid the dependency on all data modalities due to the fully connected architecture, alternatives have been proposed training VAEs on every modality separately while aligning the latent spaces^37^. This might ease the mapping of an individual to the correct position in the latent space when only some modalities are present, but the same problem arises if single features are missing within modalities.

Although the trained MOVE model was not able to significantly enhance the survival model performances of single-omics datasets, it showed some promising results when used as an imputation method. The ability to reconstruct metabolomics features despite the absence of the full metabolomics dataset was surprising, indicating that MOVE was able to learn how features across modalities were related to each other. Such a good reconstruction of the metabolomics dataset was likely due to the presence of highly related features in other datasets. We also observed a weak correlation between imputation quality and the standard deviation of the reconstructed feature. We hypothesize that this correlation stems from the fact that features with higher missingness have a lower standard deviation, and while the missingness might affect the reconstruction ability, the lower standard deviation in the data might affect the standard deviation in the reconstruction. When tracking the feature importances back from the decoder reconstruction to the encoder input, we notice similar patterns across the best-reconstructed features in different modalities. These findings link back to the observed encoder feature importances, explaining the high weights of the urine and physical measurements modalities and the influence of dataset properties on the latent representations.

In conclusion, we showed that survival models trained on a disease-independent latent representation of the data compare well against models trained on a disease-dependent subset of original features. However, we simultaneously observe strong signals from non-biological information in the dataset, which can be observed throughout our results. Factors such as data missingness, data quality, data processing and the choice of features had a deep impact on the latent representations and all downstream results. In the light of our results, we want to raise awareness for how signals coming from the actual underlying biology and the dataset properties can both have a strong impact on results when using computational methods to analyze biomedical datasets. Therefore, we see the need for further understanding and quantification of these factors’ influence on the latent representations and their impact on i) disease incidence prediction and ii) the conclusions drawn from analyzing both the survival models and the VAEs.

## Methods

### Dataset

We used data from the UK Biobank^7^, which is a dataset containing over half a million individuals from the United Kingdom and covering a wide variety of biological measurements and phenotypes. The used data was collected between 2006 and 2010, at the initial (“baseline”) visit. We exclusively used tabular data and genotype data, while the UK Biobank also contains other modalities like imaging data, electrocardiogram data and whole-genome sequencing data. Data access was granted under the application 32683 ‘Combined effect of the genetic and lifestyle determinants of metabolic syndrome on cardiometabolic risk’. All work on the data was done on the Esrum HPC cluster, provided by the Data Analytics Platform of the Novo Nordisk Foundation Center for Basic Metabolic Research.

### Data preprocessing for MOVE

We excluded individuals with 1st-degree or 2nd-degree relatedness (kinship coefficient cutoff at 0.0884). The relatedness matrix and kinship calculation were done using “KING: Kinship-based INference for Gwas”^38^ and led to the exclusion of 34,919 individuals while retaining 467,307 individuals. Additionally, we removed all individuals without genomics data, resulting in 452,026 individuals used in the analysis.

Features were selected both manually and by dimensionality reduction techniques as follows. The selected features were categorized into nine modalities: Physical measurements, metabolomics, blood cell counts, blood biochemistry, phenotype-associated genotype data (“genomics”), protein Quantitative Trait Loci (pQTL), proteomics, urine measurements and seropositivity of infectious diseases. Additionally, the features age, sex, UK Biobank assessment centre, ethnic background, fasting time, smoking status and alcohol intake frequency were added as covariates. Only data from the first UK Biobank assessment centre visit (“baseline”) was used.

Included physical measurements covered a variety of pulse, blood pressure, grip strength, height, weight and respiratory measurements. All metabolomic features were selected, except those containing percentages when there was a feature for the same metabolite describing absolute values. The selected blood biochemistry features cover a wide range of commonly used blood biomarkers and for the blood cell counts, concentrations, cell counts, and sizes of different blood cells were included. For the urine measurements and infectious disease seropositivities, all available features were included. For the proteomics dataset we used all 2,923 available proteins, which were measured by Olink^39^.

The genomic data was processed in two manners and therefore split into two separate datasets. Both datasets contained single nucleotide polymorphisms (SNP) which were encoded as 0,1,2 per individual and SNP (indicating the number of alleles deviating from the reference genome). First, the data from the Pan-UK Biobank study^40^ was used to retrieve SNPs that were significantly (p-value < 10^-16) associated with at least five traits examined in the Pan-UK Biobank study. This resulted in a matrix of (242,381 SNPs across 779 traits) containing the negative log10 of p-values. Using “DBSCAN” from the “scikit-learn” python package^41^, the SNPs were clustered for each autosome separately, and for each cluster, the SNP closest to the cluster mean was picked, in addition to all SNPs marked as outliers. Using the DBSCAN parameters eps=50 and min_samples=2 yielded 4052 SNPs, of which 3943 could be retrieved from the genotype data (**Supplementary Figure 6a**). Second, we used pQTL loci from^39^, which had -log(p-value) > 16 and were on an autosome, and retrieved the top three SNPs per protein according to their p-values. This led to 4912 SNPs, of which 3146 were found in the genotype data and 107 were shared among the two datasets (**Supplementary Figure 6b**).

A full list of included features, as well as the number of features per dataset, can be found in the online repository.

The continuous features of this dataset were standardized and the categorical features one-hot encoded using the “encode” function of MOVE, extending the 10,277 features to 24,455. Next, in order to remove the effect of the covariates, a linear regression was fitted for each continuous feature against the covariates, and the residuals were taken as the new values. Finally, the residualized features underwent standardization (calculated only on non-missing values of the training set) and all missing values were replaced by zeros. The final dataset consisted of 24,455 features across 9 modalities for 452,026 individuals.

### MOVE architecture & weights

MOVE (Multi-Omics Variational AutoEncoder) is a python package that implements multi-omics integration using a VAE architecture. Additionally, MOVE comes with added functionalities such as data preprocessing, hyperparameter tuning and feature importance calculations and can be trained with datasets containing missing data^19^. For this analysis, we exclusively used the version “developer-move-2.0”. We decided to include a single hidden layer for both the encoder and decoder while the number of neurons for the latent and hidden layers would be determined during hyperparameter tuning. MOVE also allows each dataset to have a certain weight for the loss calculation during training. We approximately set the dataset weights to the inverse of the data completeness by setting the weights for the proteomics and infectious diseases datasets to 10 and setting the weight for the metabolomics dataset to 2 while keeping the weight for all other datasets at 1.

### Quantifying clustering in latent space with k-nearest neighbors

To compare the effect of residualization on the latent representation of the VAE, we trained MOVE on three different datasets, but each based on the same randomly selected subset of 100,000 individuals. The three datasets were: i) the standardized and one-hot-encoded features including covariates, ii) the residualized features without covariates and iii) the same as i) but setting the MOVE weights of the covariates for the loss calculation to 0. Given the three trained models, we then calculated the mean accuracy over all samples when predicting the labels of the categorical covariates with the “KNeighborsClassifier” from the “scikit-learn” package^41^. This was done for different values of k between k=10 (local clustering) and k=99,999 (convergence to majority class).

### Train and test data splits

The sampled individuals were randomly split into 90% train and 10% test sets. This was automatically done by MOVE and the same train and test splits were further used for the subsequent survival models. After training the largest MOVE model on 360,000 of 400,000 sampled individuals, the remaining 52,026 individuals were kept for the imputation analysis and the single-omics experiments, again separated into 90% train and 10% test (**Supplementary Figure 7**). This separation into two cohorts, each with their own train and test sets, prevented data leakage when using a pre-trained MOVE model.

### MOVE hyperparameter tuning & training

To select the set of hyperparameters to be used with MOVE for further analysis, we tuned for the number of neurons in the hidden layers, the number of neurons in the bottleneck layer, and the weight of the Kullback-Leibler Divergence (KLD). For these models, 100000 randomly sampled individuals were used (**Supplementary Figure 8**). After choosing a suitable set of hyperparameters (KLD weight: 0.001, 512 hidden neurons, 128 latent neurons), we trained a bigger model on 400,000 individuals, while the remaining 52,026 individuals were kept for further analysis. 10% of the samples were held out as the test set and used to calculate reconstruction metrics, resulting in an actual training set of size 360,000 and a test set of size 40,000. Training was done for 100 epochs using the Adam optimizer with a learning rate of 0.0001.

### MOVE feature importance analysis

To analyze MOVE, we used the “IntegratedGradients” function of the “Captum” package^42^, which is built to interpret “PyTorch”-based neural networks. We calculated the attributions for encoder and decoder separately, using a randomly selected subset of 5,000 samples due to computational cost. For each input feature for the encoder and decoder, we took the mean of the absolute attributions over all samples, which resulted in two tables of size (24,455 features x 128 latent dimensions) that were used for subsequent analysis. The encoder feature importances were calculated by summing up all means of absolute attributions of an input feature across all latent neurons and were used as a measure of feature importance for the latent representation. For the feature importance for a certain reconstructed or imputed feature, we matrix-multiplied the decoder importance vector (128 dimensions) for the reconstructed feature with the encoder importance matrix, resulting in a vector of size (24,455 input features).

### Survival models, disease selection and disease-specific data preprocessing

To predict disease incidence, we used four types of survival models: CoxPHFitter and LogNormalAFTFitter from the “lifelines” package^43^, as well as two XGBoost models from the “xgboost” package^31^, using the objectives “survival:cox” and “survival:aft” respectively. We selected

15 diseases considering the following factors (in no specific order): Number of reported cases, availability of self-reported cases at baseline, variety of ICD-10 chapters and clear definition with ICD-10 codes. The corresponding set of ICD-10 codes was retrieved from the HDR UK Phenotype Library^44^ for each disease and can be found in the online repository, while the incidence distributions can be seen in **Supplementary Figure 9**. For each disease, individuals who met a previously defined set of criteria^25^ were excluded from both the training and test sets. In addition to that, individuals which died or were diagnosed earlier than 180 days after their first UK Biobank (baseline) visit were removed from the disease-specific dataset. This also included self-reported diagnoses at the baseline visit. The censoring time for the survival model was calculated as the difference between the in-hospital diagnosis date and the UK Biobank baseline visit date for the individuals diagnosed within 10 years, and as the minimum time from the baseline visit to either death, loss of follow-up, or the 10-year censoring period otherwise. Finally, we trained all four model types on 2 different input datasets, which were the latent representation of the previously trained MOVE model and a disease-specific lasso-selected subset of the residualized input features.

### Survival model training and evaluation

The survival models were fitted on the training set (90%) and evaluated on the test set (10%) as defined by the prior MOVE training. For exact model parameters, we refer to the published code. The trained models were then evaluated on the test set by calculating the C-index (higher values indicating better model performance) with the “concordance_index_censored” function of the package “scikit-survival”^45^. As this function calculates the C-index based on hazards and the C-index calculation is based on ranking, we calculated the C-index on the negative predicted survival time if a survival model was not able to return a hazard prediction.

### Result variability analysis

To understand the effect of training both the VAE and the survival models, we ran 10 rounds of VAE training and subsequent survival model training, or 10 rounds of survival models on the same latent space, respectively. In the first case, we fixed the seed for everything but the VAE training, whereas in the second case, we fixed the seed for everything but the survival model training. In both cases, we used the same hyperparameters and dataset weights as for the big MOVE model (on 400,000 samples) and used 100,000 randomly selected samples for these models. We chose to restrict the survival models to a subset of five diseases and only trained survival models on the latent space.

### Survival model feature importances

In order to analyze the trained survival models, we calculated feature importances for the XGBoost models using Shapley additive explanations from the “shap” package^23^, where we averaged the absolute values per feature over all training samples. For the Cox proportional hazards and accelerated failure time models, we retrieved model coefficients to better understand which features had the most influence on disease incidence prediction. For the models based on the latent representation, this resulted in a matrix of shape (15 diseases x 128 latent dimensions) per model type (**Supplementary Figure 4**). To calculate the correlation between C-index and feature importances, we summed the feature importances per disease and model type, resulting in four correlation values. To link the survival model feature importances to the input features of the encoder, we matrix-multiplied the mean absolute encoder importances with the mean absolute survival model importances to create a vector of size (24,455 input features) per disease and model type.

### Mapping single-omics datasets to the latent space

To examine the ability of the trained MOVE model to integrate samples from a single omics dataset into the existing latent space, therefore possibly enhancing the disease prediction, we used the remaining 52,026 individuals which were not used in the MOVE model to test this hypothesis. We looped over all datasets, always keeping data from one while setting the features of all others to zero, and mapped those samples to the latent space using the pre-trained encoder. We then trained and evaluated survival models on those latent representations, again randomly spitting the data into 90% training samples and 10% test samples. For comparison, we also trained and evaluated survival models on a corresponding lasso-selected subset of residualized input data (only containing values for one dataset) and, if feasible (<= 100 features), on the whole residualized input data.

### Imputing missing values with trained MOVE

We used the trained MOVE model to understand to what extent it can be used to impute missing values. We first examined how well the imputation works if of the existing values, 2,000 were removed randomly from each feature simultaneously. In a second scenario, we removed all the data for all samples and features from a single dataset and repeated this for every dataset. For both scenarios, we then calculated the Pearson correlation between the removed and reconstructed values, as well as the standard deviation of each reconstructed feature. For this analysis, we exclusively used the 52,026 samples that were not used for MOVE training or testing beforehand.

## Supporting information

Supplementary Infomation

## Acknowledgements

C.R., J.F.G., R.H.M., M.P.A., H.W., K.R. and S.R. were supported by the Novo Nordisk Foundation (grant NNF23SA0084103). J.F.G. was supported by a research grant from the Danish Cardiovascular Academy, which is funded by the Novo Nordisk Foundation, grant number NNF20SA0067242 and The Danish Heart Foundation. M.P.A was funded by the Novo Nordisk Foundation Copenhagen Bioscience Ph.D. Program grant No. NNF22SA0078229.

## Author contributions

Conceptualization: S.R. Formal Analysis: C.R., J.F.G. Investigation: C.R. Resources: S.R. Software: C.R., R.H.M., H.W. Supervision: S.R. Validation: C.R. Visualization: C.R. Writing – original draft: C.R. Writing – review & editing: S.R., J.F.G., M.P.A, R.H.M., H.W., K.R.

## Data availability

For access to the UK Biobank data, researchers must apply for approval (https://www.ukbiobank.ac.uk/). Data from the Pan-UK Biobank study used for the SNP clustering is publicly available on https://pan.ukbb.broadinstitute.org/.

## Code availability

The code, parameters and lists of used features and ICD-10 codes can be retrieved from https://github.com/RasmussenLab/move_ukbb.

## References

1. Julkunen, H. et al. Atlas of plasma NMR biomarkers for health and disease in 118,461 individuals from the UK Biobank. Nat Commun 14, 604 (2023).

2. Deng, Y.-T. et al. Atlas of the plasma proteome in health and disease in 53,026 adults. Cell (2024).

3. Cai, L. et al. Causal associations between cardiorespiratory fitness and type 2 diabetes. Nat Commun 14, 3904 (2023).

4. Pandey, A. et al. Biomarker-based risk prediction of incident heart failure in pre-diabetes and diabetes. Heart Fail 9, 215–223 (2021).

5. Suzuki, K. et al. Genetic drivers of heterogeneity in type 2 diabetes pathophysiology. Nature 627, 347–357 (2024).

6. Shields, B. M. et al. Patient stratification for determining optimal second-line and third-line therapy for type 2 diabetes: the TriMaster study. Nat Med 29, 376–383 (2023).

7. Sudlow, C. et al. UK biobank: an open access resource for identifying the causes of a wide range of complex diseases of middle and old age. PLoS Med 12, e1001779 (2015).

8. Investigators, A. The “All of Us” research program. New England Journal of Medicine 381, 668–676 (2019).

9. Lonsdale, J. et al. The genotype-tissue expression (GTEx) project. Nat Genet 45, 580–585 (2013).

10. Wörheide, M. A., Krumsiek, J., Kastenmüller, G. & Arnold, M. Multi-omics integration in biomedical research–A metabolomics-centric review. Anal Chim Acta 1141, 144–162 (2021).

11. Picard, M., Scott-Boyer, M.-P., Bodein, A., Périn, O. & Droit, A. Integration strategies of multi-omics data for machine learning analysis. Comput Struct Biotechnol J 19, 3735–3746 (2021).

12. Beesley, L. J. et al. The emerging landscape of health research based on biobanks linked to electronic health records: Existing resources, statistical challenges, and potential opportunities. Stat Med 39, 773–800 (2020).

13. Kingma, D. P. Auto-encoding variational bayes. arXiv preprint arXiv:1312.6114 (2013).

14. Sieliwonczyk, E. et al. Unsupervised electrocardiogram feature extraction using deep learning empowers discovery of genetic and phenotypic determinants of cardiac electrophysiology. Eur Heart J 45, ehae666–3441 (2024).

15. Puyol-Antón, E. et al. Assessing the impact of blood pressure on cardiac function using interpretable biomarkers and variational autoencoders. in International Workshop on Statistical Atlases and Computational Models of the Heart 22–30 (Springer, 2019).

16. Way, G. P. & Greene, C. S. Extracting a biologically relevant latent space from cancer transcriptomes with variational autoencoders. in PACIFIC SYMPOSIUM on BIOCOMPUTING 2018: Proceedings of the Pacific Symposium 80–91 (2018).

17. Gomari, D. P. et al. Variational autoencoders learn transferrable representations of metabolomics data. Commun Biol 5, 645 (2022).

18. Hira, M. T. et al. Integrated multi-omics analysis of ovarian cancer using variational autoencoders. Sci Rep 11, 6265 (2021).

19. Allesøe, R. L. et al. Discovery of drug–omics associations in type 2 diabetes with generative deep-learning models. Nat Biotechnol 41, 399–408 (2023).

20. Rasheed, K. et al. Explainable, trustworthy, and ethical machine learning for healthcare: A survey. Comput Biol Med 149, 106043 (2022).

21. Tasin, I., Nabil, T. U., Islam, S. & Khan, R. Diabetes prediction using machine learning and explainable AI techniques. Healthc Technol Lett 10, 1–10 (2023).

22. Raihan, M. J., Khan, M. A.-M., Kee, S.-H. & Nahid, A.-A. Detection of the chronic kidney disease using XGBoost classifier and explaining the influence of the attributes on the model using SHAP. Sci Rep 13, 6263 (2023).

23. Lundberg, S. A unified approach to interpreting model predictions. arXiv preprint arXiv:1705.07874 (2017).

24. Cox, D. R. Regression models and life-tables. Journal of the Royal Statistical Society: Series B (Methodological) 34, 187–202 (1972).

25. Carrasco-Zanini, J. et al. Proteomic signatures improve risk prediction for common and rare diseases. Nat Med 30, 2489–2498 (2024).

26. You, J. et al. Plasma proteomic profiles predict individual future health risk. Nat Commun 14, 7817 (2023).

27. Buergel, T. et al. Metabolomic profiles predict individual multidisease outcomes. Nat Med 28, 2309–2320 (2022).

28. Wiegrebe, S., Kopper, P., Sonabend, R., Bischl, B. & Bender, A. Deep learning for survival analysis: a review. Artif Intell Rev 57, 65 (2024).

29. Salerno, S. & Li, Y. High-dimensional survival analysis: Methods and applications. Annu Rev Stat Appl 10, 25–49 (2023).

30. Wei, L.-J. The accelerated failure time model: a useful alternative to the Cox regression model in survival analysis. Stat Med 11, 1871–1879 (1992).

31. Chen, T. & Guestrin, C. Xgboost: A scalable tree boosting system. in Proceedings of the 22nd acm sigkdd international conference on knowledge discovery and data mining 785–794 (2016).

32. Committee, P. & others. 2. Diagnosis and Classification of Diabetes: Standards of Care in Diabetes-2024. Diabetes Care 47, S20–S42 (2024).

33. Jawaid, I. & Rajesh, S. Hyperparathyroidism (primary) NICE guideline: diagnosis, assessment, and initial management. The British Journal of General Practice 70, 362 (2020).

34. Stensrud, M. J. & Hernán, M. A. Why test for proportional hazards? JAMA 323, 1401–1402 (2020).

35. Sjölander, A. & Dickman, P. W. Why test for proportional hazards—or any other model assumptions? Am J Epidemiol 193, 926–927 (2024).

36. Uno, H., Cai, T., Pencina, M. J., D’Agostino, R. B. & Wei, L.-J. On the C-statistics for evaluating overall adequacy of risk prediction procedures with censored survival data. Stat Med 30, 1105–1117 (2011).

37. Kutuzova, S., Igel, C., Nielsen, M. & McCloskey, D. Bi-modal variational autoencoders for metabolite identification using tandem mass spectrometry. bioRxiv 2021–2028 (2021).

38. Manichaikul, A. et al. Robust relationship inference in genome-wide association studies. Bioinformatics 26, 2867–2873 (2010).

39. Sun, B. B. et al. Plasma proteomic associations with genetics and health in the UK Biobank. Nature 622, 329–338 (2023).

40. Karczewski, K. J. et al. Pan-UK Biobank GWAS improves discovery, analysis of genetic architecture, and resolution into ancestry-enriched effects. MedRxiv 2023–2024 (2024).

41. Pedregosa, F. et al. Scikit-learn: Machine learning in Python. the Journal of machine Learning research 12, 2825–2830 (2011).

42. Kokhlikyan, N. et al. Captum: A unified and generic model interpretability library for pytorch. arXiv preprint arXiv:2009.07896 (2020).

43. Davidson-Pilon, C. lifelines: survival analysis in Python. J Open Source Softw 4, 1317 (2019).

44. Thayer, D. S. et al. Creating a next-generation phenotype library: the health data research UK Phenotype Library. JAMIA Open 7, (2024).

45. Pölsterl, S. scikit-survival: A Library for Time-to-Event Analysis Built on Top of scikit-learn. Journal of Machine Learning Research 21, 1–6 (2020).

