## Supplementary Infomation for "Unsupervised learning of multi-omics data enables disease risk prediction in the UK Biobank"

**Supplementary Information**

**A. Result tables for single-omics datasets**

Models: CPH = Cox Proportional Hazards. AFT = Accelerated Failure Time. XGB = XGBoost.
Input: Latent = Latent representation as input. Lasso = Lasso-selected feature subset as input. Full = All input features as input (only done if no more than 100 features)

| Condition | **Latent CPH** | **Lasso CPH** | **Full XGBCPH** | **Latent XGBCPH** | **Lasso XGBCPH** | **Full XGBAFT** | **Latent XGBAFT** | **Lasso XGBAFT** | **Latent AFT** | **Lasso AFT** |
|---|---|---|---|---|---|---|---|---|---|---|
| **anaemia** | 0.541 | 0.512 | 0.000 | 0.532 | 0.000 | 0.000 | **0.543** | 0.509 | 0.543 | 0.511 |
| **angina pectoris** | **0.512** | 0.465 | 0.000 | 0.501 | 0.000 | 0.000 | 0.507 | 0.452 | 0.512 | 0.464 |
| **copd** | 0.510 | 0.493 | 0.000 | 0.513 | 0.000 | 0.000 | **0.516** | 0.513 | 0.514 | 0.492 |
| **depression** | 0.503 | 0.510 | 0.000 | 0.519 | 0.000 | 0.000 | **0.520** | 0.506 | 0.503 | 0.511 |
| **primary hypertension** | 0.509 | 0.518 | 0.000 | 0.501 | 0.000 | 0.000 | 0.495 | 0.518 | 0.509 | **0.519** |
| **renal failure** | 0.453 | 0.455 | 0.000 | 0.465 | 0.000 | 0.000 | 0.464 | **0.484** | 0.456 | 0.457 |
| **type 2 diabetes mellitus** | 0.525 | 0.564 | 0.000 | 0.535 | 0.000 | 0.000 | 0.542 | 0.560 | 0.523 | **0.567** |

**Supplementary Table 1**: C-indices based on genomics dataset, filtered for at least 100 incident test cases. Highest C-index per row is highlighted.

| Condition | **Latent CPH** | **Lasso CPH** | **Full XGBCPH** | **Latent XGBCPH** | **Lasso XGBCPH** | **Full XGBAFT** | **Latent XGBAFT** | **Lasso XGBAFT** | **Latent AFT** | **Lasso AFT** |
|---|---|---|---|---|---|---|---|---|---|---|
| **anaemia** | 0.518 | 0.489 | 0.000 | **0.527** | 0.000 | 0.000 | 0.525 | 0.490 | 0.516 | 0.488 |
| **angina pectoris** | 0.498 | 0.499 | 0.000 | 0.480 | 0.000 | 0.000 | 0.499 | 0.491 | **0.505** | 0.498 |
| **copd** | 0.487 | 0.509 | 0.000 | 0.511 | 0.000 | 0.000 | **0.520** | 0.500 | 0.488 | 0.510 |
| **depression** | 0.470 | 0.484 | 0.000 | **0.514** | 0.000 | 0.000 | 0.502 | 0.494 | 0.464 | 0.487 |
| **primary hypertension** | 0.487 | **0.512** | 0.000 | 0.508 | 0.000 | 0.000 | 0.507 | 0.510 | 0.487 | 0.511 |
| **renal failure** | **0.520** | 0.510 | 0.000 | 0.501 | 0.000 | 0.000 | 0.498 | 0.510 | 0.519 | 0.510 |
| **type 2 diabetes mellitus** | **0.524** | 0.521 | 0.000 | 0.513 | 0.000 | 0.000 | 0.517 | 0.511 | 0.524 | 0.523 |

**Supplementary Table 2**: C-indices based on pQTL dataset, filtered for at least 100 incident test cases. Highest C-index per row is highlighted.

| Condition | **Latent CPH** | **Lasso CPH** | **Full XGBCPH** | **Latent XGBCPH** | **Lasso XGBCPH** | **Full XGBAFT** | **Latent XGBAFT** | **Lasso XGBAFT** | **Latent AFT** | **Lasso AFT** |
|---|---|---|---|---|---|---|---|---|---|---|
| **anaemia** | 0.617 | 0.623 | 0.651 | 0.638 | 0.651 | 0.651 | 0.640 | **0.651** | 0.618 | 0.623 |
| **angina pectoris** | 0.570 | 0.606 | 0.612 | 0.594 | 0.615 | **0.623** | 0.600 | 0.622 | 0.569 | 0.606 |
| **copd** | 0.610 | 0.595 | 0.712 | 0.639 | 0.000 | **0.713** | 0.642 | 0.703 | 0.618 | 0.596 |
| **depression** | 0.592 | 0.591 | **0.602** | 0.563 | 0.599 | 0.600 | 0.564 | 0.600 | 0.594 | 0.591 |
| **primary hypertension** | 0.630 | 0.620 | **0.668** | 0.652 | 0.000 | 0.667 | 0.652 | 0.667 | 0.632 | 0.621 |
| **renal failure** | 0.715 | 0.710 | 0.770 | 0.737 | 0.000 | **0.772** | 0.735 | 0.764 | 0.738 | 0.721 |

| | | | | | | | | | | |
|---|---|---|---|---|---|---|---|---|---|---|
| **type 2 diabetes mellitus** | 0.851 | 0.837 | 0.882 | 0.858 | 0.882 | **0.883** | 0.861 | 0.883 | 0.851 | 0.850 |

**Supplementary Table 3**: C-indices based on blood biochemistry dataset, filtered for at least 100 incident test cases. Highest C-index per row is highlighted.

| Condition | **Latent CPH** | **Lasso CPH** | **Full XGBCPH** | **Latent XGBCPH** | **Lasso XGBCPH** | **Full XGBAFT** | **Latent XGBAFT** | **Lasso XGBAFT** | **Latent AFT** | **Lasso AFT** |
|---|---|---|---|---|---|---|---|---|---|---|
| **anaemia** | 0.676 | 0.674 | 0.678 | 0.672 | 0.677 | **0.679** | 0.678 | 0.678 | 0.677 | 0.676 |
| **angina pectoris** | 0.607 | 0.569 | 0.623 | 0.594 | 0.614 | **0.626** | 0.596 | 0.613 | 0.602 | 0.572 |
| **copd** | 0.618 | 0.631 | 0.676 | 0.648 | 0.000 | 0.681 | 0.647 | **0.684** | 0.619 | 0.633 |
| **depression** | 0.561 | 0.548 | **0.599** | 0.542 | 0.586 | 0.582 | 0.535 | 0.575 | 0.561 | 0.546 |
| **primary hypertension** | 0.583 | 0.592 | 0.605 | 0.583 | 0.000 | 0.604 | 0.583 | **0.605** | 0.584 | 0.590 |
| **renal failure** | 0.628 | 0.597 | **0.633** | 0.614 | 0.000 | 0.632 | 0.621 | 0.632 | 0.623 | 0.604 |
| **type 2 diabetes mellitus** | 0.691 | 0.682 | 0.718 | 0.692 | 0.715 | **0.720** | 0.689 | 0.718 | 0.692 | 0.688 |

**Supplementary Table 4**: C-indices based on blood cell count dataset, filtered for at least 100 incident test cases. Highest C-index per row is highlighted.

| Condition | **Latent CPH** | **Lasso CPH** | **Full XGBCPH** | **Latent XGBCPH** | **Lasso XGBCPH** | **Full XGBAFT** | **Latent XGBAFT** | **Lasso XGBAFT** | **Latent AFT** | **Lasso AFT** |
|---|---|---|---|---|---|---|---|---|---|---|
| **anaemia** | 0.613 | 0.627 | 0.000 | 0.648 | 0.000 | 0.000 | **0.649** | 0.640 | 0.618 | 0.634 |
| **primary hypertension** | 0.636 | 0.640 | 0.000 | 0.633 | 0.000 | 0.000 | 0.634 | 0.633 | 0.640 | **0.641** |
| **renal failure** | 0.710 | 0.717 | 0.000 | 0.721 | 0.000 | 0.000 | 0.725 | **0.727** | 0.711 | 0.719 |
| **type 2 diabetes mellitus** | 0.803 | 0.831 | 0.000 | 0.821 | 0.000 | 0.000 | 0.824 | **0.856** | 0.826 | 0.842 |

**Supplementary Table 5**: C-indices based on metabolomics dataset, filtered for at least 100 incident test cases. Highest C-index per row is highlighted.

| Condition | **Latent CPH** | **Lasso CPH** | **Full XGBCPH** | **Latent XGBCPH** | **Lasso XGBCPH** | **Full XGBAFT** | **Latent XGBAFT** | **Lasso XGBAFT** | **Latent AFT** | **Lasso AFT** |
|---|---|---|---|---|---|---|---|---|---|---|
| **anaemia** | 0.596 | 0.601 | 0.602 | 0.593 | 0.602 | 0.607 | 0.600 | **0.608** | 0.598 | 0.600 |
| **angina pectoris** | 0.575 | **0.634** | 0.612 | 0.614 | 0.633 | 0.619 | 0.617 | 0.632 | 0.585 | 0.634 |
| **copd** | 0.734 | 0.723 | **0.799** | 0.769 | 0.000 | 0.799 | 0.769 | 0.799 | 0.749 | 0.735 |
| **depression** | 0.575 | 0.568 | **0.596** | 0.566 | 0.588 | 0.584 | 0.564 | 0.581 | 0.569 | 0.570 |
| **primary hypertension** | 0.678 | 0.668 | **0.689** | 0.686 | 0.000 | 0.689 | 0.687 | 0.689 | 0.681 | 0.669 |
| **renal failure** | 0.667 | 0.646 | 0.672 | **0.680** | 0.000 | 0.674 | 0.678 | 0.674 | 0.668 | 0.646 |
| **type 2 diabetes mellitus** | 0.748 | 0.753 | 0.756 | 0.752 | 0.758 | 0.755 | 0.753 | **0.758** | 0.749 | 0.754 |

**Supplementary Table 6**: C-indices based on physical measurements dataset, filtered for at least 100 incident test cases. Highest C-index per row is highlighted.

| Condition | **Latent CPH** | **Lasso CPH** | **Full XGBCPH** | **Latent XGBCPH** | **Lasso XGBCPH** | **Full XGBAFT** | **Latent XGBAFT** | **Lasso XGBAFT** | **Latent AFT** | **Lasso AFT** |
|---|---|---|---|---|---|---|---|---|---|---|
| **anaemia** | 0.540 | 0.509 | **0.544** | 0.507 | **0.544** | 0.544 | 0.506 | 0.544 | 0.531 | 0.509 |
| **angina pectoris** | **0.591** | 0.510 | 0.573 | 0.582 | 0.573 | 0.572 | 0.578 | 0.572 | 0.580 | 0.511 |
| **copd** | 0.577 | 0.522 | 0.581 | 0.532 | 0.000 | **0.582** | 0.526 | **0.582** | 0.578 | 0.518 |
| **depression** | 0.459 | 0.569 | 0.517 | 0.513 | 0.482 | 0.525 | 0.504 | 0.487 | 0.471 | **0.574** |
| **primary hypertension** | 0.580 | 0.548 | 0.603 | 0.597 | 0.000 | **0.606** | 0.597 | 0.606 | 0.585 | 0.546 |
| **renal failure** | 0.572 | 0.569 | 0.615 | 0.596 | 0.000 | 0.615 | 0.597 | **0.615** | 0.578 | 0.569 |
| **type 2 diabetes mellitus** | 0.664 | 0.563 | 0.663 | 0.639 | 0.663 | 0.667 | 0.637 | **0.667** | 0.658 | 0.562 |

**Supplementary Table 7**: C-indices based on urine dataset, filtered for at least 100 incident test cases. Highest C-index per row is highlighted.

### B. Supplementary figures

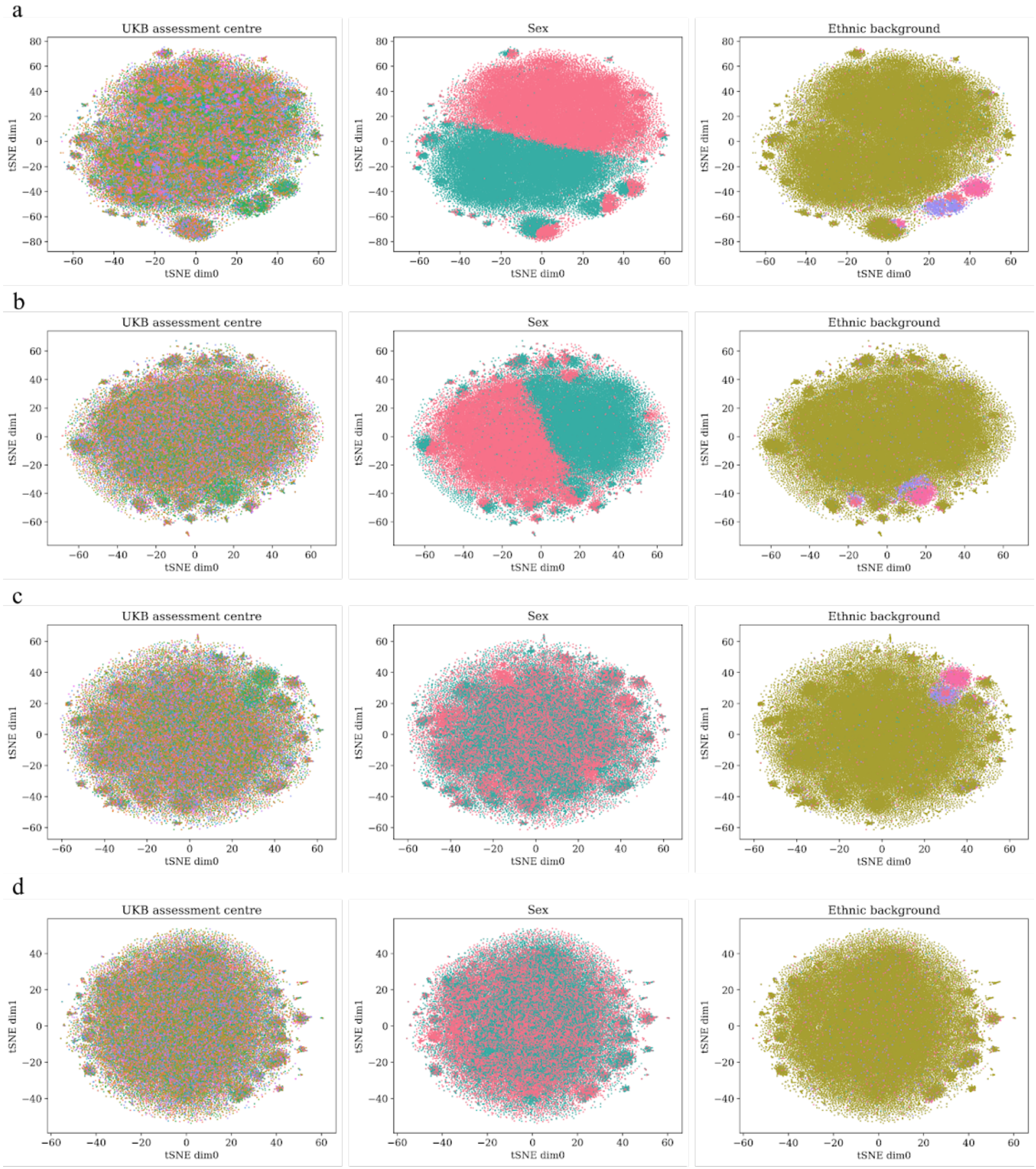


**Supplementary Figure 1: Clustering of latent space from different datasets.** Plots show t-distributed Stochastic Neighbour Embedding (t-SNE)[1] of latent representation, colored by three different categorical covariates (UK Biobank assessment centre, Sex, Ethnic background, from left to right). 100,000 randomly sampled individuals were used. **a**) Training MOVE on a non-residualized dataset with categorical genotype data (including the covariates). **b**) Training MOVE on a non-residualized dataset with categorical genotype data (including the covariates) but setting the covariate weights to 0. **c**) Training MOVE on a residualized dataset with non-residualized categorical genotype data. **d**) Training MOVE on a residualized dataset with continuous and residualized genotype data.

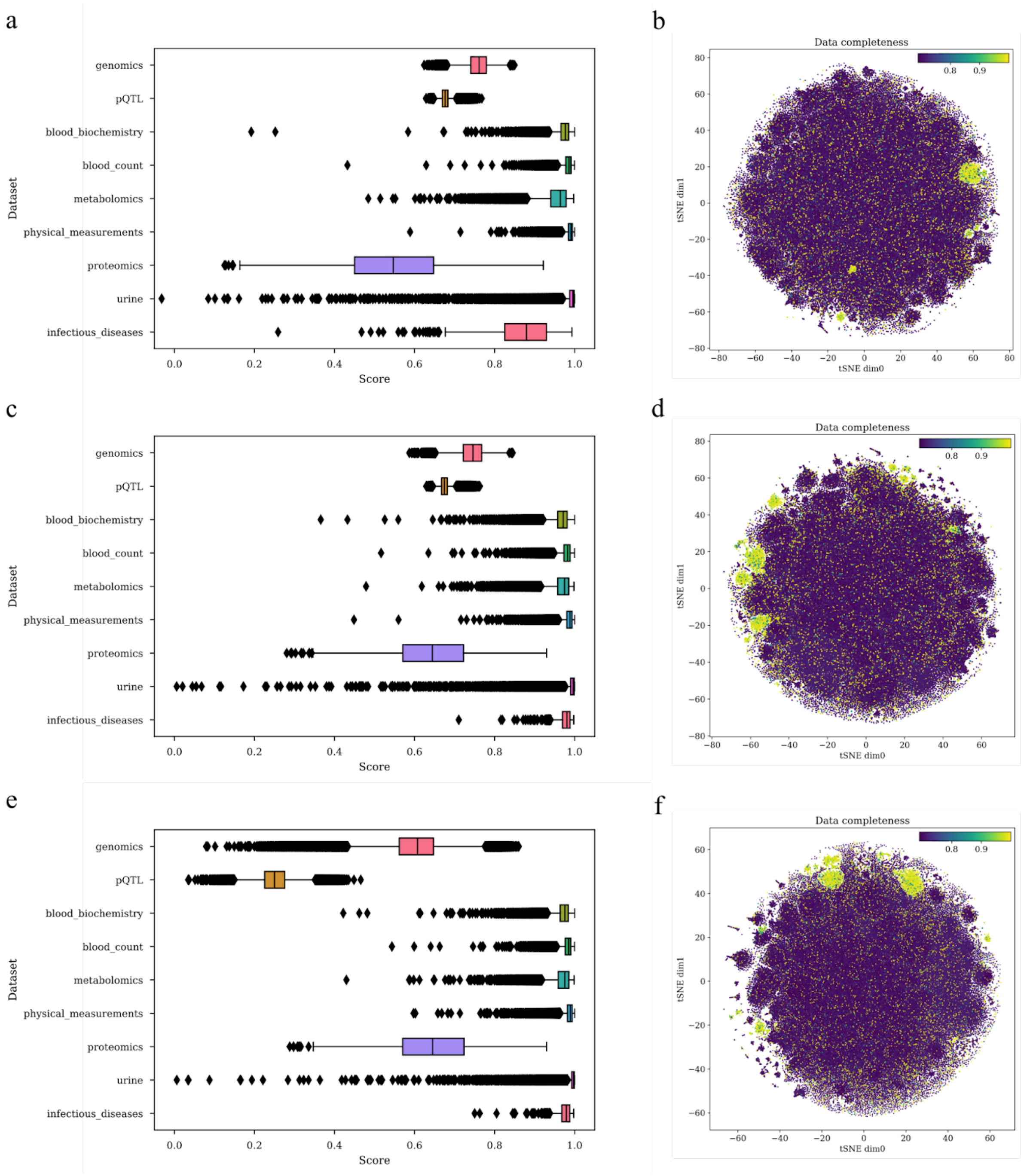


**Supplementary Figure 2: Effect of dataset weight adjustment.** Comparison of reconstruction metrics (left; cosine similarity for continuous modalities and accuracy for categorical modalities) and clustering in the t-SNE-reduced latent space by data completeness (right), using 400,000 randomly sampled individuals. **a-b**) before adjusting weights and using categorical genomics and pQTL data, **c-d**) after adjusting weights and using categorical genomics and pQTL data, **e-f**) after adjusting weights and using continuous genomics and pQTL data.

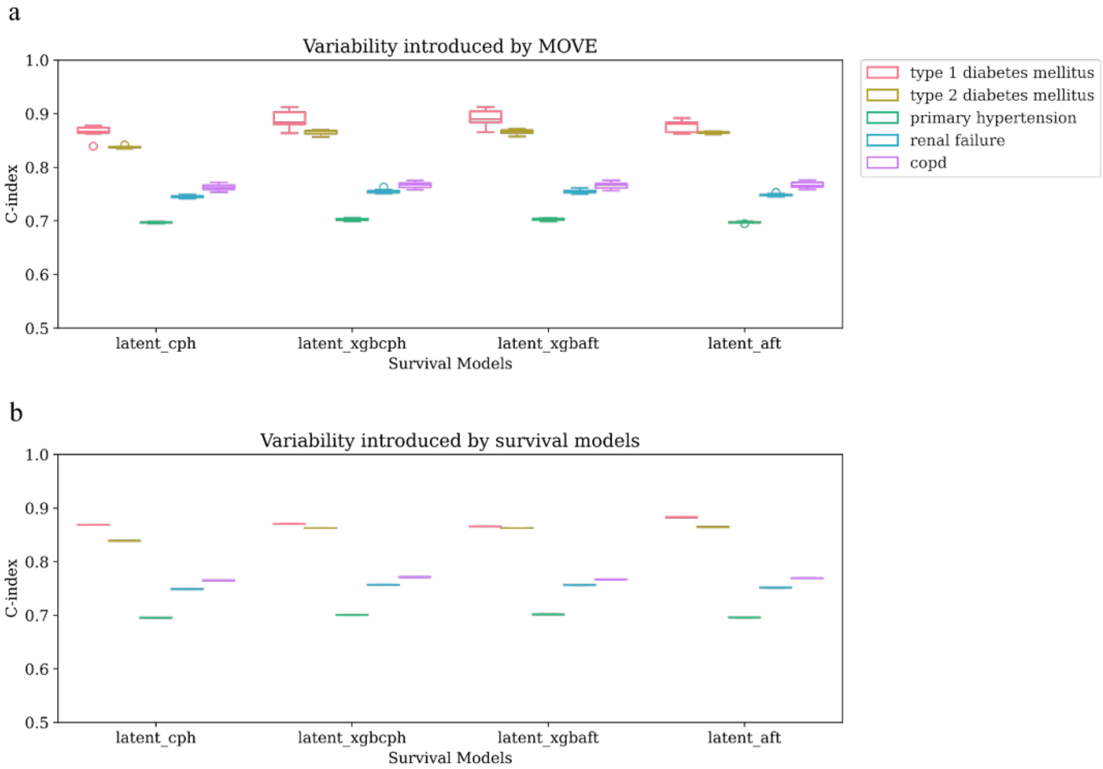


**Supplementary Figure 3: Result variability introduced by MOVE and survival models.** For all latent models and five diseases, models were trained and evaluated 10 times. **a**) Variability by MOVE while keeping seed of the survival model constant. **b**) Variability by survival models based on the same latent space.

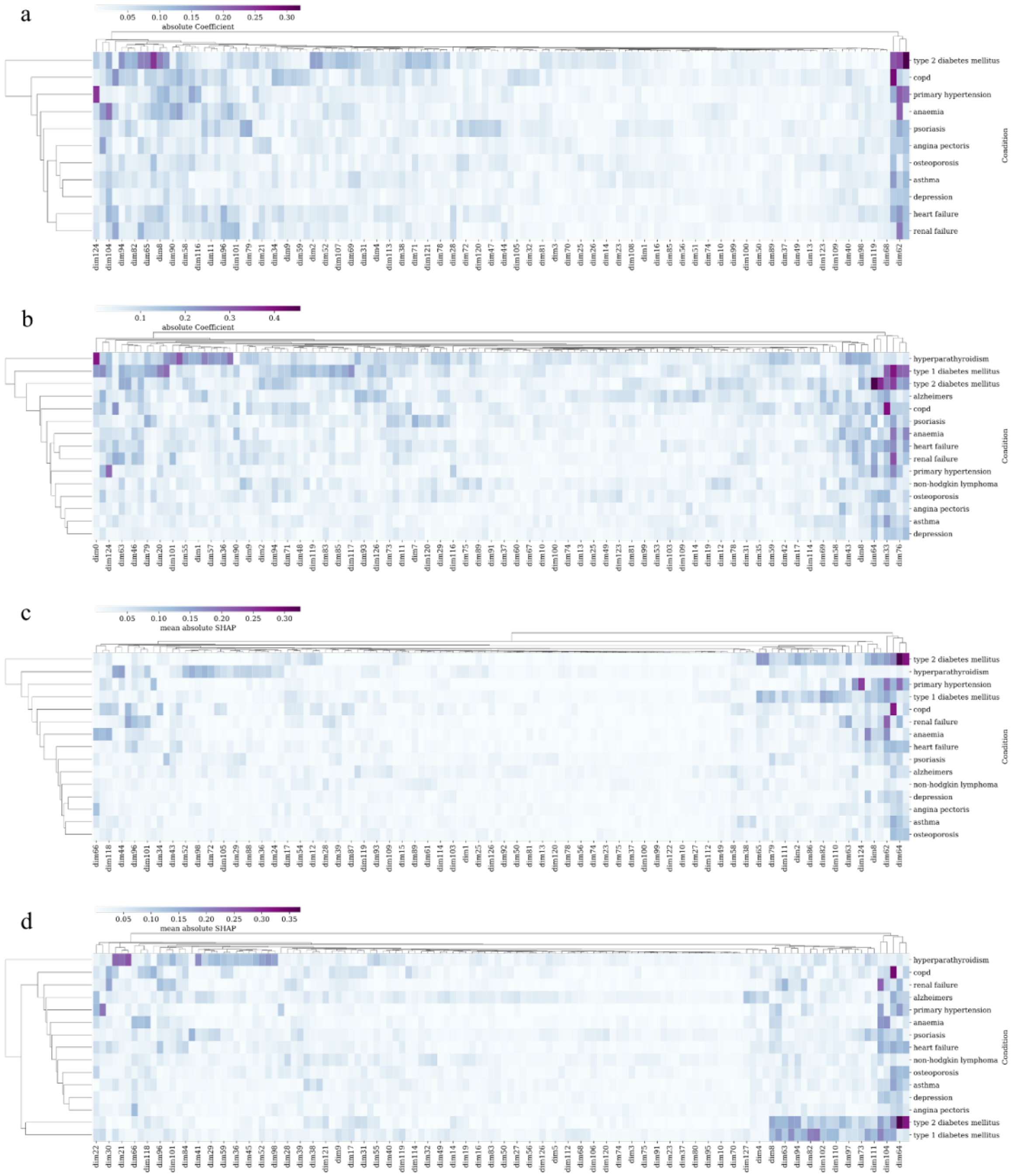


**Supplementary Figure 4: Feature importances per model and disease.** All models were trained on the latent representation. **a**) Absolute coefficients of Accelerated Failure Time models. **b**) Absolute coefficients of Cox Proportional Hazards models. **c**) Mean absolute SHAP values for XGBoost models with AFT-loss. **d**) Mean absolute SHAP values for XGBoost models with Cox-loss.

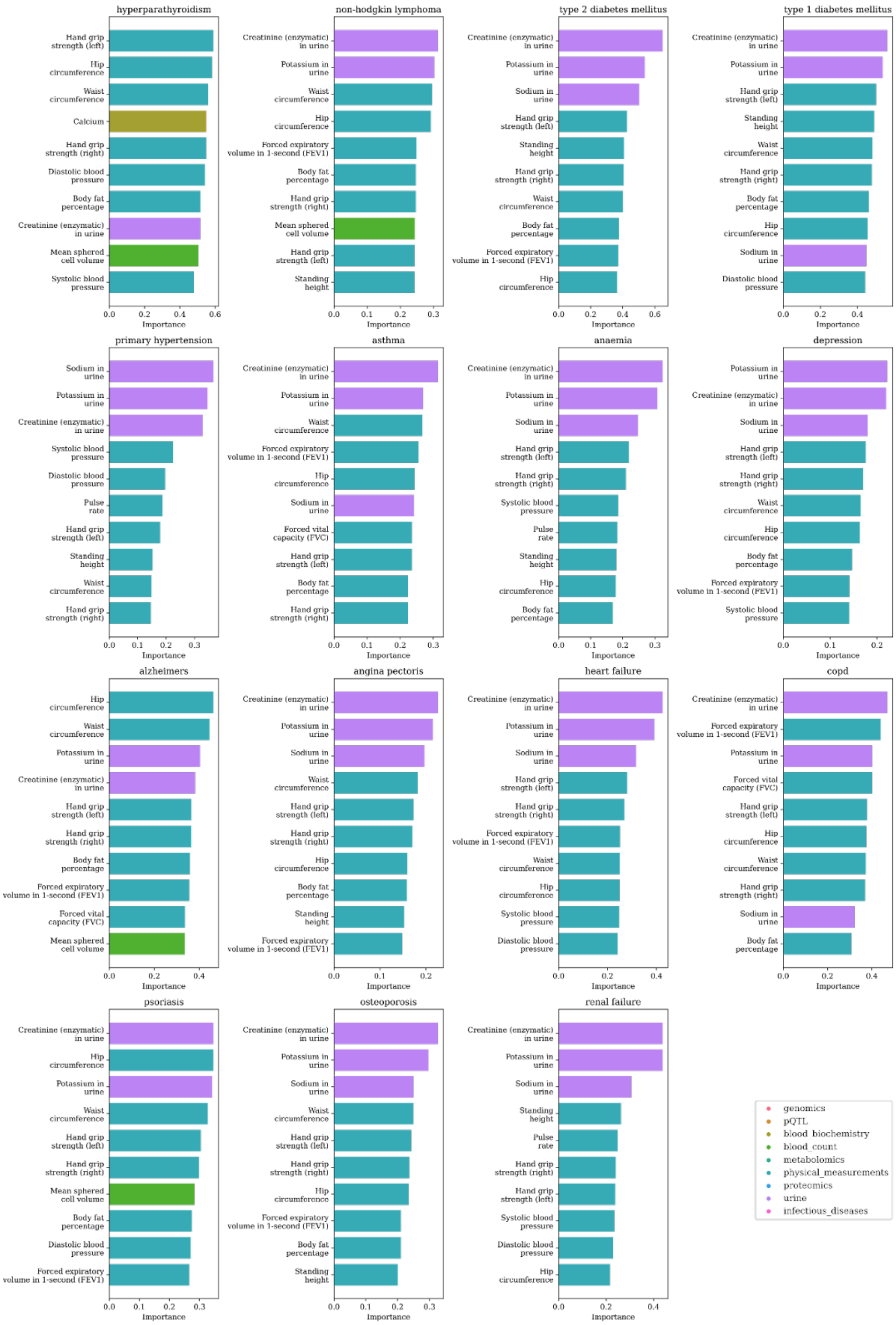

hyperparathyroidism
Hand grip strength (left)
Hip circumference
Waist circumference
Calcium
Hand grip strength (right)
Diastolic blood pressure
Body fat percentage
Creatinine (enzymatic) in urine
Mean sphered cell volume
Systolic blood pressure
Importance
non-hodgkin lymphoma
Creatinine (enzymatic) in urine
Potassium in urine
Waist circumference
Hip circumference
Forced expiratory volume in 1-second (FEV1)
Body fat percentage
Hand grip strength (right)
Mean sphered cell volume
Hand grip strength (left)
Standing height
Importance
type 2 diabetes mellitus
Creatinine (enzymatic) in urine
Potassium in urine
Sodium in urine
Hand grip strength (left)
Standing height
Hand grip strength (right)
Waist circumference
Body fat percentage
Forced expiratory volume in 1-second (FEV1)
Hip circumference
Importance
type 1 diabetes mellitus
Creatinine (enzymatic) in urine
Potassium in urine
Hand grip strength (left)
Standing height
Waist circumference
Hand grip strength (right)
Body fat percentage
Hip circumference
Sodium in urine
Diastolic blood pressure
Importance
primary hypertension
Sodium in urine
Potassium in urine
Creatinine (enzymatic) in urine
Systolic blood pressure
Diastolic blood pressure
Pulse rate
Hand grip strength (left)
Standing height
Waist circumference
Hand grip strength (right)
Importance
asthma
Creatinine (enzymatic) in urine
Potassium in urine
Waist circumference
Forced expiratory volume in 1-second (FEV1)
Hip circumference
Sodium in urine
Forced vital capacity (FVC)
Hand grip strength (left)
Body fat percentage
Hand grip strength (right)
Importance
anaemia
Creatinine (enzymatic) in urine
Potassium in urine
Sodium in urine
Hand grip strength (left)
Hand grip strength (right)
Systolic blood pressure
Pulse rate
Standing height
Hip circumference
Body fat percentage
Importance
depression
Potassium in urine
Creatinine (enzymatic) in urine
Sodium in urine
Hand grip strength (left)
Hand grip strength (right)
Waist circumference
Hip circumference
Body fat percentage
Forced expiratory volume in 1-second (FEV1)
Systolic blood pressure
Importance
alzheimers
Hip circumference
Waist circumference
Potassium in urine
Creatinine (enzymatic) in urine
Hand grip strength (left)
Hand grip strength (right)
Body fat percentage
Forced expiratory volume in 1-second (FEV1)
Forced vital capacity (FVC)
Mean sphered cell volume
Importance
angina pectoris
Creatinine (enzymatic) in urine
Potassium in urine
Sodium in urine
Waist circumference
Hand grip strength (left)
Hand grip strength (right)
Hip circumference
Body fat percentage
Standing height
Forced expiratory volume in 1-second (FEV1)
Importance
heart failure
Creatinine (enzymatic) in urine
Potassium in urine
Sodium in urine
Hand grip strength (left)
Hand grip strength (right)
Forced expiratory volume in 1-second (FEV1)
Waist circumference
Hip circumference
Systolic blood pressure
Diastolic blood pressure
Importance
copd
Creatinine (enzymatic) in urine
Forced expiratory volume in 1-second (FEV1)
Potassium in urine
Forced vital capacity (FVC)
Hand grip strength (left)
Hip circumference
Waist circumference
Hand grip strength (right)
Sodium in urine
Body fat percentage
Importance
psoriasis
Creatinine (enzymatic) in urine
Hip circumference
Potassium in urine
Waist circumference
Hand grip strength (left)
Hand grip strength (right)
Mean sphered cell volume
Body fat percentage
Diastolic blood pressure
Forced expiratory volume in 1-second (FEV1)
Importance
osteoporosis
Creatinine (enzymatic) in urine
Potassium in urine
Sodium in urine
Waist circumference
Hand grip strength (left)
Hand grip strength (right)
Hip circumference
Forced expiratory volume in 1-second (FEV1)
Body fat percentage
Standing height
Importance
renal failure
Creatinine (enzymatic) in urine
Potassium in urine
Sodium in urine
Standing height
Pulse rate
Hand grip strength (right)
Hand grip strength (left)
Systolic blood pressure
Diastolic blood pressure
Hip circumference
Importance
genomics
pQTL
blood_biochemistry
blood_count
metabolomics
physical_measurements
proteomics
urine
infectious_diseases

**Supplementary Figure 5: Feature importances for disease predictions from latent representation.** Top 10 features by mean absolute SHAP-values per predicted disease when using the XGBoost model with Cox-Loss. Colored by modality and displayed by predicted disease.

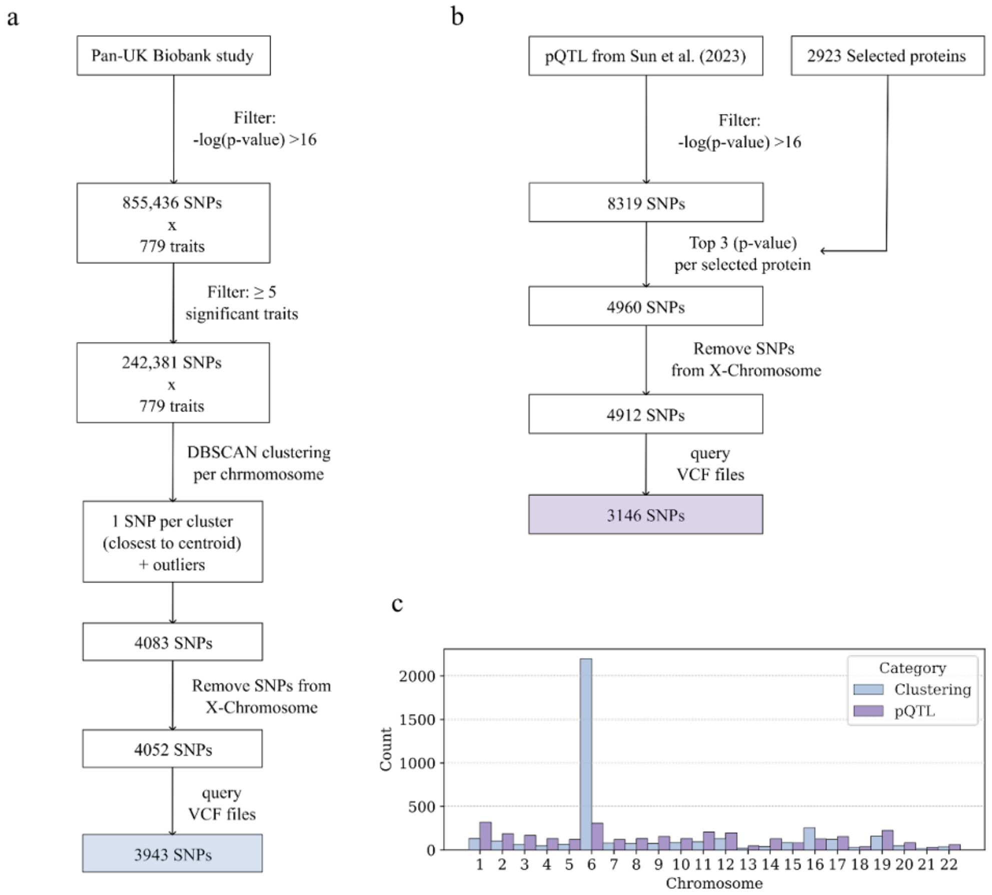


**Supplementary Figure 6: Overview of single-nucleotide polymorphism (SNP) selection for the genomics and pQTL datasets.** a) Steps to select SNPs with a clustering approach. b) Steps to select SNPs by taking the top three protein quantitative trait loci (pQTL) per protein. c) Histogram displaying number of selected SNPs per autosome and dataset.

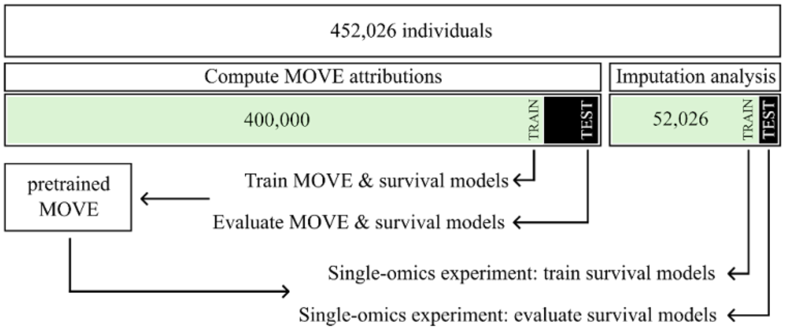


**Supplementary Figure 7: Overview of train / test split.** Out of 452,026 individuals, 400,000 were used to train (90%) and evaluate (10%) the biggest MOVE model and subsequent survival models. The pre-trained MOVE model was later used to map the remaining 52,026 individuals to the latent space while doing the "single-omics" experiments. Of these 52,026 individuals, 90% were used to train survival models while the remaining 10% were used as test samples. Furthermore, 5,000 of the 400,000 individuals were randomly selected to compute the attributions of MOVE, while the 52,026 individuals were used to analyze the performance of MOVE as an imputation method.

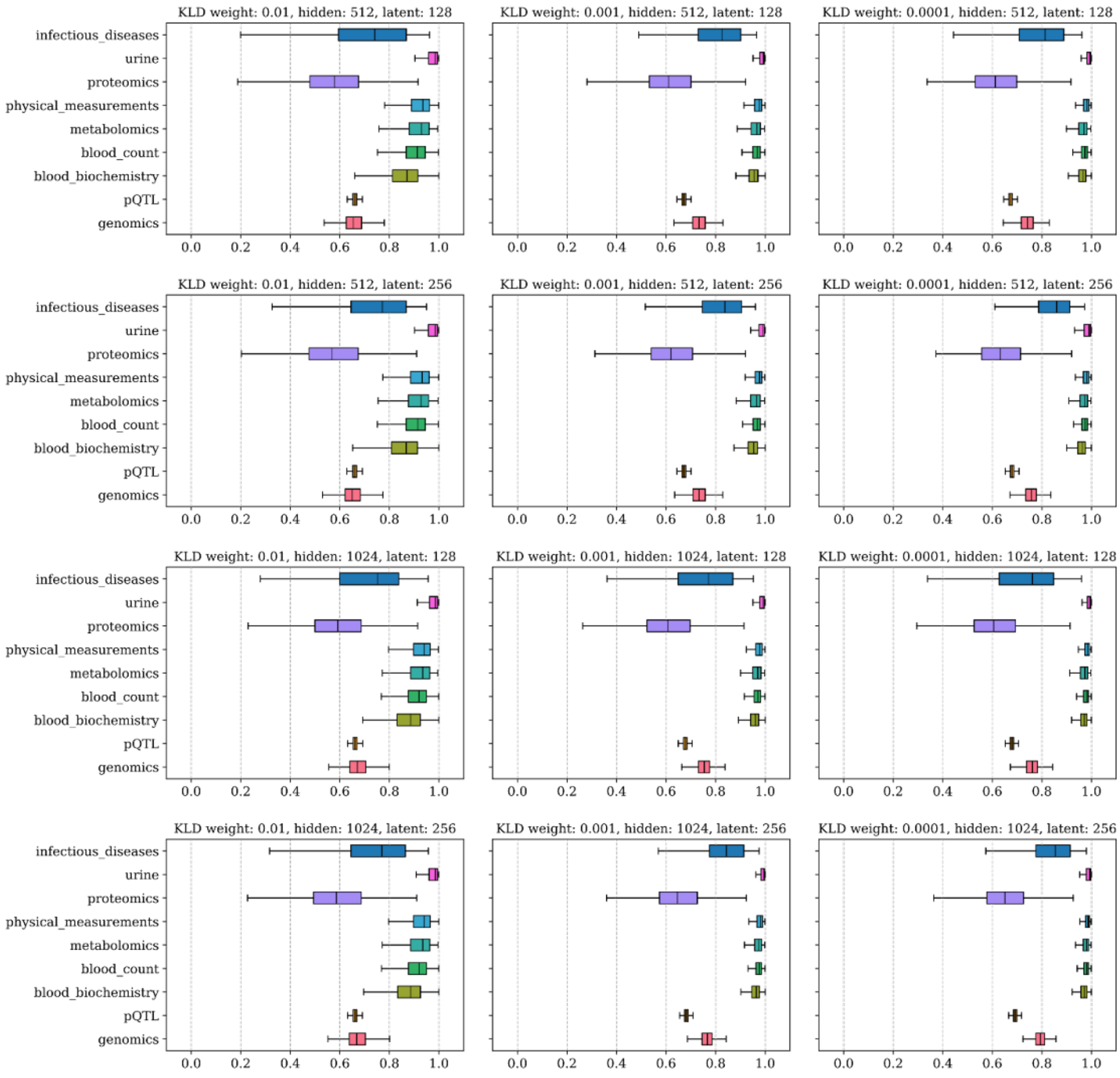


**Supplementary Figure 8: MOVE hyperparameter tuning results.** Parameters included in tuning are: Kullback-Leibler divergence (KLD) weight, number of hidden neurons, number of latent neurons. Boxplots show cosine similarity for the continuous modalities and accuracy for the categorical modalities (genomics and pQTL) between input features and reconstruction across all individuals per dataset.

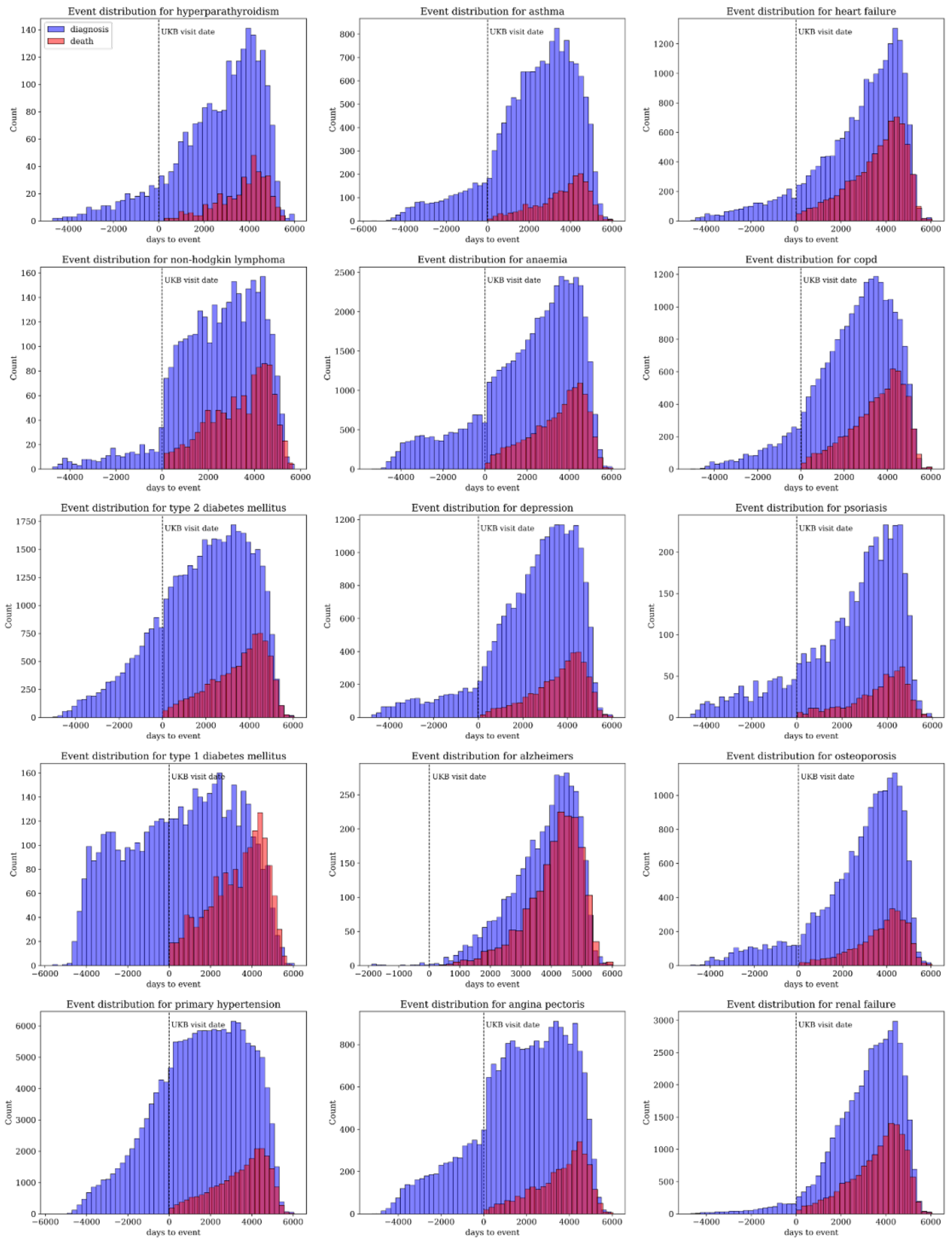


**Supplementary Figure 9:** Distribution of incidences for predicted diseases and deaths of the corresponding individuals. Vertical line indicates time point of UK Biobank baseline visit. Death cause is not constrained to disease.

**References for Supplementary Section**

1. Van der Maaten, L. & Hinton, G. Visualizing data using t-SNE. *Journal of machine learning research* 9, (2008).
